# Isolating the sources of heterogeneity in nanoparticle-cell interactions

**DOI:** 10.1101/817569

**Authors:** Stuart T Johnston, Matthew Faria, Edmund J Crampin

**Affiliations:** Systems Biology Laboratory, School of Mathematics and Statistics, and Department of Biomedical Engineering, University of Melbourne, Parkville, Victoria 3010, Australia; ARC Centre of Excellence in Convergent Bio-Nano Science and Technology, Melbourne School of Engineering, University of Melbourne, Parkville, Victoria 3010, Australia; School of Medicine, Faculty of Medicine Dentistry and Health Sciences, University of Melbourne, Parkville, Victoria 3010, Australia

**Keywords:** Cell heterogeneity, nanoparticles, mathematical modeling, nanoparticle-cell in-teractions, cell cycle

## Abstract

Nanoparticles have the potential to enhance therapeutic success and reduce toxicity-based treatment side effects via the targeted delivery of drugs to cells. This delivery relies on complex interactions between numerous biological, chemical and physical processes. The intertwined nature of these processes has thus far hindered attempts to understand their individual impact. Variation in experimental data, such as the number of nanoparticles inside each cell, further inhibits understanding. Here we present a mathematical framework that is capable of examining the impact of individual processes during nanoparticle delivery. We demonstrate that variation in experimental nanoparticle uptake data can be explained by three factors: random nanoparticle motion; variation in nanoparticle-cell interactions; and variation in the maximum nanoparticle uptake per cell. Without all three factors, the experimental data cannot be explained. This work provides insight into biological mechanisms that cause heterogeneous responses to treatment, and enables precise identification of treatment-resistant cell subpopulations.

## 1 Introduction

Elucidating how individual biological and physical processes dictate the successful cellular uptake of nanoparticles is crucial for future developments in areas such as nanomedicine and nanotoxicology (1, 2). Untangling the role of a particular process requires a detailed understanding of the complex marriage of transport phenomena, physicochemical nanoparticle characteristics and biological behavior that govern nanoparticle-cell interactions (1–4). It is well established that the physicochemical properties of a nanoparticle, such as size, shape and surface charge, impact nanoparticle uptake (1, 2, 4, 5). The exact influence of these properties is unclear, as the impact is obscured by both the influence of transport phenomena, such as sedimentation, diffusion and aggregation, and cell-type specific interactions between cells and nanoparticles (4, 6–8). The inherent variation in cell characteristics within a population further obscures the roles of individual processes, and results in heterogeneous experimental data (9–13).

Heterogeneity in experimental data may imply that commonly-reported population-averaged measures do not accurately reflect the underlying biology (14–16). For example, consider a nanoparticle-cell association assay, where the average number of nanoparticles associated with a cell is measured after exposure to a particular concentration of nanoparticles (7). This measure, referred to as nanoparticle dose, can be used as a proxy for the effectiveness of a putative treatment (17). Effective treatment of disease via drug-loaded nanoparticles may require universal cellular uptake within a population (12), such that all cells interact with the drug. Reporting only the average number of associated nanoparticles does not distinguish between the influence of stochastic processes, where all cells are identical but associate with nanoparticles at random, and the presence of distinct subpopulations of cells that interact differently with nanoparticles due to fundamental differences in biology (Fig. 1). Certain cell subpopulations may not associate with nanoparticles, or associate with nanoparticles at an inhibited rate (12). Even if such cell subpopulations are rare, they can nevertheless have a substantial impact on disease progression (14, 18, 19). Identifying whether a cell population does, in fact, contain heterogeneity in relevant cell characteristics or whether variation in experimental data is merely a by-product of the stochastic nature of nanoparticle transport is therefore critical for therapeutic success (9–13). Furthermore, understanding how heterogeneity in cell characteristics, such as receptor numbers or vesicle formation rates, manifests itself in commonly-measured experimental data is crucial for isolating and quantifying sources of heterogeneity.

**Figure 1:**
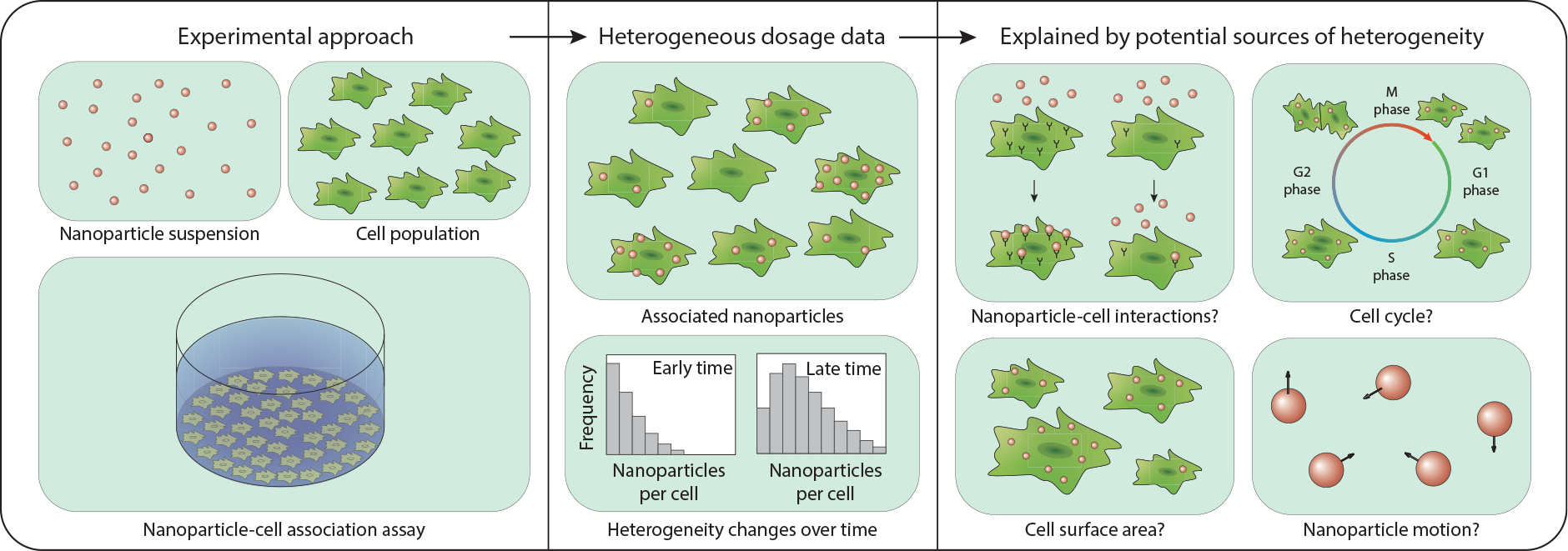
Schematic highlighting the experimental approach that gives rise to a heterogeneous dosage distribution with potential sources of heterogeneity. Nanoparticle-cell association assays result in heterogeneous dosage distributions, which may be explained by heterogeneity in (i) nanoparticle-cell interactions; (ii) the cell cycle; (iii) cell surface area or; (iv) stochastic nanoparticle motion.

Nanoparticle motion is inherently stochastic due to the fundamental length scales involved in the transport process (3, 11). As such, the measured nanoparticle dose per cell will be distributed according to the transport properties, as well as any potential heterogeneity in cell characteristics. Without careful consideration of the contribution of the stochastic nature of transport to the dosage distribution, heterogeneity in cell characteristics can be misidentified or incorrectly estimated. Mathematical models of nanoparticle motion are effective at isolating the contribution of nanoparticle transport to dosage from biological interactions (3, 5–8). However, such models describe the average nanoparticle behavior and dose, and are not suitable for predicting dosage distributions or cell heterogeneity. Statistical approaches allow for the quantification of heterogeneity from experimental data (9, 10, 13), but do not provide mechanistic understanding about how heterogeneity in experimental data arises from heterogeneity in multiple cell characteristics. Further, as we will demonstrate, conclusions obtained from previous statistical approaches are incapable of explaining heterogeneity observed in our time course experiments.

## Results

Here we develop and introduce a model of individual nanoparticle behavior that is capable of describing and predicting cell heterogeneity. This modeling framework mimics experimental conditions while providing detail at both an individual particle and individual cell level. The standard experimental approach for analyzing nanoparticle-cell interactions is an adherent cell culture association assay (6). In an association assay, a cell population seeded on a culture dish is incubated in media containing a nanoparticle suspension (Fig. 1) (6). The nanoparticles undergo transport through the fluid via a combination of sedimentation and diffusion, and ultimately arrive at the cell-media interface (Fig. 2C,E-G) (3, 6–8). Nanoparticles bind to receptors on the cell surface and are internalized via various endocytic processes (4). The evolution of the number of nanoparticles associated with each cell is measured to provide time course information on the dose (Fig. 2D). It is difficult to distinguish between nanoparticles that are internalized by a cell or are merely bound to the cell surface, and hence we take the standard approach of using the number of associated nanoparticles as a proxy for dose (7, 20). Due to the ubiquitous use of association experiments to investigate nanoparticle efficacy, we calibrate the geometry and conditions in our modeling framework to an association assay.

**Figure 2:**
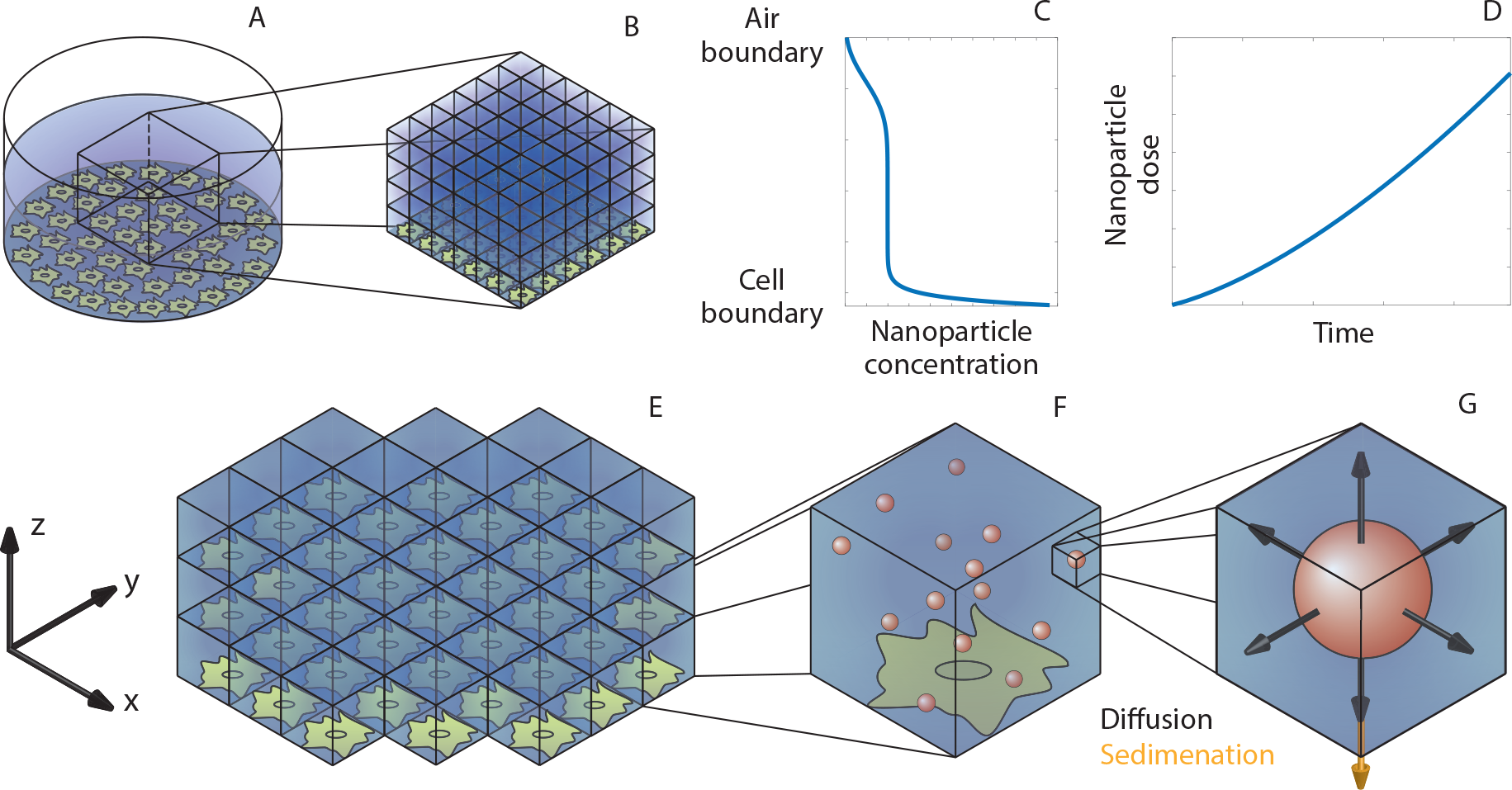
Experimental and model geometry. (**A**) Standard experimental geometry for an *in vitro* adherent cell culture association assay. (**B**) Representative geometry for the voxel-based modeling framework. (**C**) Typical nanoparticle concentration as a function of depth due to sedimentation and diffusion of nanoparticles. (**D**) Typical nanoparticle association curve. (E) Media-cell boundary in the modeling framework highlighting (**F**) nanoparticle locations within a voxel and (**G**) the contributions of random motion (diffusion) and directed motion (sedimentation) to nanoparticle transport.

We implement a voxel-based framework, where the experimental domain is discretized into cube-shaped subdomains known as voxels (Fig. 2A-B). We model the number of nanoparticles within each voxel, which evolves with time due to stochastic transitions of nanoparticles between voxels. The transition rates correspond to the combined rates of sedimentation and diffusion (21). Transition events are sampled via a spatial stochastic simulation algorithm, a modified form of the well-established Gillespie’s algorithm (22). To replicate experimental conditions, there is no transition of nanoparticles through the top of the domain, corresponding to the air-media interface. At the cell-media interface, the transition rate of nanoparticles from the media into the cell monolayer corresponds to the cell carrying capacity kinetics derived by Faria *et al.* (7). These kinetics have been demonstrated to be the most suitable kinetics for describing association assays for a wide range of nanoparticle-cell combinations (7). The kinetics rely on two parameters: a *nanoparticle-cell affinity* parameter, which represents the rate of interaction between a nanoparticle and a cell, and; a *cell carrying capacity* parameter, which is the maximum number of nanoparticles that can associate with a cell (7). The equivalence of the transition rate used here to the cell carrying capacity kinetics of Faria *et al.* (7) and an efficient method to calculate the average behavior in the voxel-based model are derived in the Supplementary Information (21, 23) and Methods section, respectively.

### Stochastic motion does not account for all observed variation

Our modeling framework describes both individual nanoparticles and individual cells and, therefore, we are able to explicitly measure the number of nanoparticles associated with each cell. As nanoparticle motion is stochastic, the model output will be a distribution of nanoparticles per cell. This distribution can be readily compared with experimental data, as flow cytometry techniques can be used to measure the number of nanoparticles associated with each cell for an entire cell population (24–26).

Experimentally, even when all cells have identical characteristics, the number of nanoparticles per cell will be heterogeneous due to the inherent stochasticity of nanoparticle motion. This dosage distribution is dependent on the association regime of the cells. For example, if the cells are in the linear association regime, where the number of nanoparticles per cell is significantly lower than the carrying capacity, then the dosage distribution will be Poisson distributed with an arrival rate equivalent to the average association rate of nanoparticles with the cell layer. However, this may not be the case when the nanoparticle dose is close to the maximum number of nanoparticles per cell.

To determine the influence of stochastic transport on the nanoparticle dosage distribution, we perform simulations using our modeling framework and calculate the number of nanoparticles associated with each cell. Here we first consider idealized conditions where all cells in an experiment have identical characteristics. These simulations are calibrated to match three different experiments: the first is performed with 1032nm PMA_SH_ capsule nanoparticles and RAW264.7 cells, the second is performed with 282nm PMA_SH_ coreshell nanoparticles and HeLa cells, and the third is performed with 150nm PMA_SH_ coreshell nanoparticles and RAW264.7 cells (7, 27). As noted previously, the dose obtained from the model is only Poisson distributed in the linear association regime (Supplementary Information, Fig. S1). If the number of nanoparticles per cell approaches the carrying capacity, the predicted dosage distribution is not well described by the Poisson distribution. Comparing the dosage distribution obtained from the model with the experimental data, we observe that the experimental data is overdisperse compared to the model predictions (Supplementary Information, Fig. S1). This indicates that the cell populations exhibit heterogeneity, and is consistent with previous observations (9–13).

### Heterogeneity is time dependent

Having established that the experimental data is not consistent with a homogeneous cell population, we next consider implementing heterogeneity in our modeling framework. A natural choice is to allow each cell in the model to have a nanoparticle-cell affinity parameter that is sampled from a probability distribution. This represents variation in the biological processes that dictate nanoparticle association, such as the number of receptors or vesicle formation rates (10). Lognormal distributions are prevalent throughout biology, and arise from multiplicative sources of variability (3, 12, 28). As such, here we make the assumption that affinity is lognormally distributed. Note that other distributions could be considered, and while this would affect the final dosage distribution, the analysis techniques remain the same.

As each cell now has an individual affinity parameter, and the association rate for each cell is proportional to the affinity, the number of nanoparticles per cell will follow a Poisson-lognormal distribution (29). Critically, this allows us to determine the relative contributions of stochastic nanoparticle motion and cell heterogeneity to the variation in the dosage distribution. The applicability of the Poisson-lognormal distribution relies on the assumption that the number of nanoparticles associated with a cell is independent of other cells; that is, competition between cells for nanoparticles is minimal. This is appropriate provided that association occurs sufficiently slowly compared to nanoparticle transport, as is the case for the nanoparticle-cell combinations considered here. There is only a single free parameter in the distribution, the standard deviation, as the mean of the Poisson-lognormal distribution must correspond to the mean number of associated nanoparticles. We refer to this standard deviation as the “apparent heterogeneity.”

The apparent heterogeneity is a key concept for the work presented here. As the dosage distribution is described by the Poisson-lognormal distribution, the heterogeneity present in the data is captured via the measure of spread in the Poisson-lognormal distribution: the standard deviation (apparent heterogeneity). It is important to note that this is not the “true” heterogeneity in one (or more) of the cell characteristics. Rather, the apparent heterogeneity represents how the “true” heterogeneity manifests itself in the experimental dosage distribution. As the nanoparticle dose can be indicative of therapeutic success, it is therefore necessary to understand how the apparent heterogeneity arises from the “true” heterogeneity in the cell characteristics.

To examine the apparent heterogeneity present in the experimental data, we fit the Poisson-lognormal distribution to the experimental data at each measured time point. In Fig. 3 we present both the distribution fit and the evolution of the apparent heterogeneity. Fig 3D-F shows that the Poisson-lognormal describes the data well for all three experiments, indicating that the assumption of lognormally-distributed cell characteristics is appropriate. Notably, the apparent heterogeneity changes with time (Fig 3A-C). For each nanoparticle-cell pair, the apparent heterogeneity decreases rapidly at early time before beginning to plateau towards the final experimental observation. This observation suggests that previous investigations into heterogeneity in nanoparticle-cell interactions, where the heterogeneity is assumed to be constant (10), do not represent a complete picture due to the time-dependent nature of nanoparticle cell interactions. Therefore, we next seek to determine how the time dependence of apparent heterogeneity arises as a consequence of interactions between nanoparticles and various cell characteristics.

**Figure 3:**
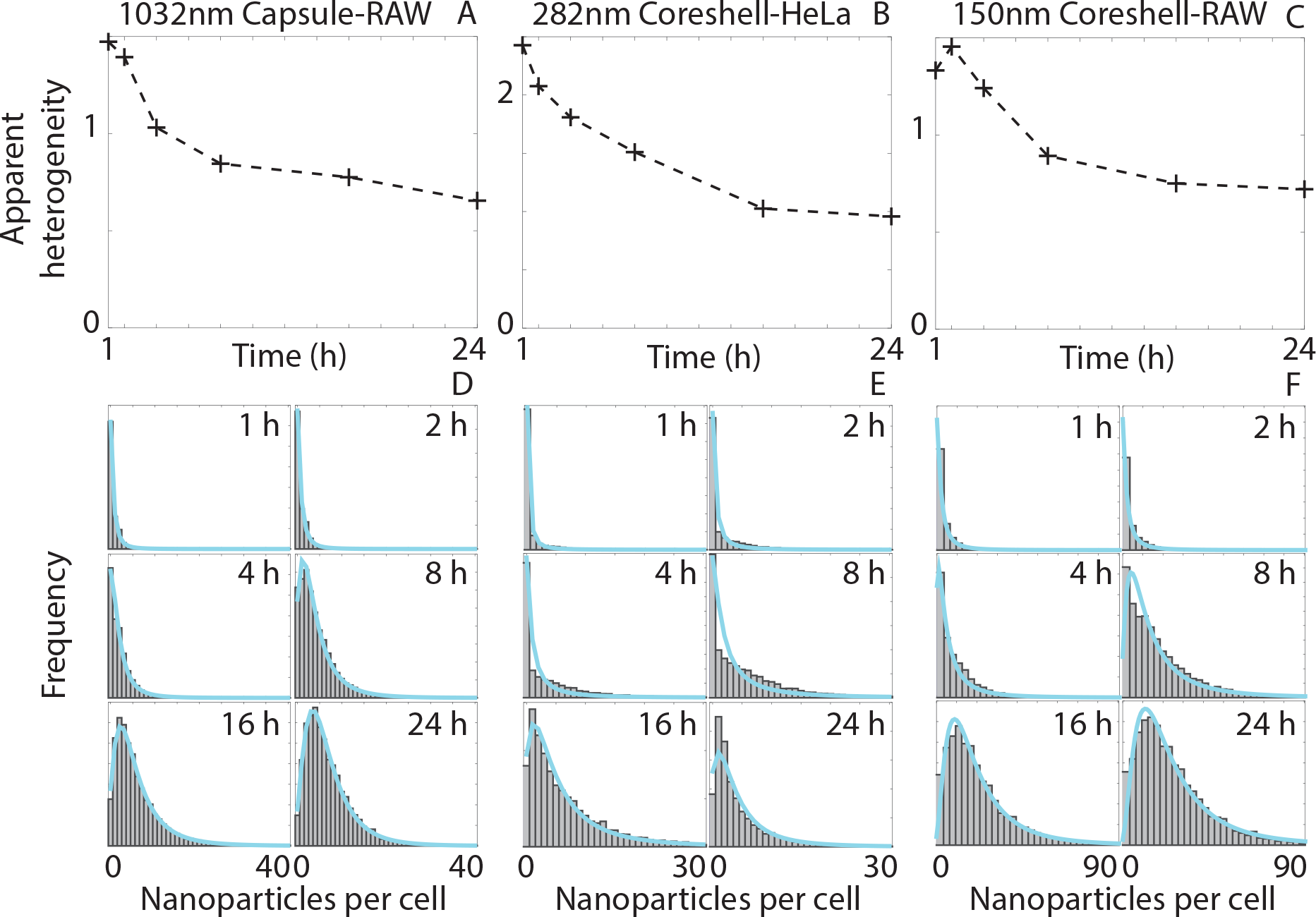
Heterogeneity appears to change with time. (**A-C**) Evolution of apparent heterogeneity obtained from experimental data for three nanoparticle-cell pairs. (**D-F**) Dosage distributions for the three nanoparticle-cell pairs after 1, 2, 4, 8, 16 and 24 hours. The cyan line corresponds to the Poisson-lognormal distribution that best fits the dosage distribution.

### Cell size distribution does not account for time-dependent heterogeneity

To determine whether the experimental apparent heterogeneity arises solely from heterogeneous nanoparticle-cell association, as suggested previously (10), we introduce cell heterogeneity into our modeling framework via the affinity parameter. A potential explanation for heterogeneous affinity is the heterogeneity in cell surface area due to the cell cycle (10, 30, 31). Specifically, we assume that a cell with higher surface area is more likely to interact and associate with nanoparticles (10). We calibrate a lognormal distribution to the square of small-angle light scattering intensity obtained via flow cytometry for both HeLA and RAW264.7 cells to estimate the cell surface area heterogeneity (32). This heterogeneity is used to create the lognormal distribution for the affinity parameter.

We perform simulations representing the three previously-described experiments, now incorporating heterogeneous nanoparticle-cell affinity, and present the results in Fig. 4A-F. To obtain robust estimates of the apparent heterogeneity, we use an efficient approach that provides the average nanoparticle dose for each cell, and fit the Poisson-lognormal distribution to this dosage data. This approach provides results that are consistent with the average output of the voxel-based model (Supplementary Information, Fig. S3.), and avoids fluctuations in the apparent heterogeneity due to the stochastic nature of the voxel-based model.

**Figure 4:**
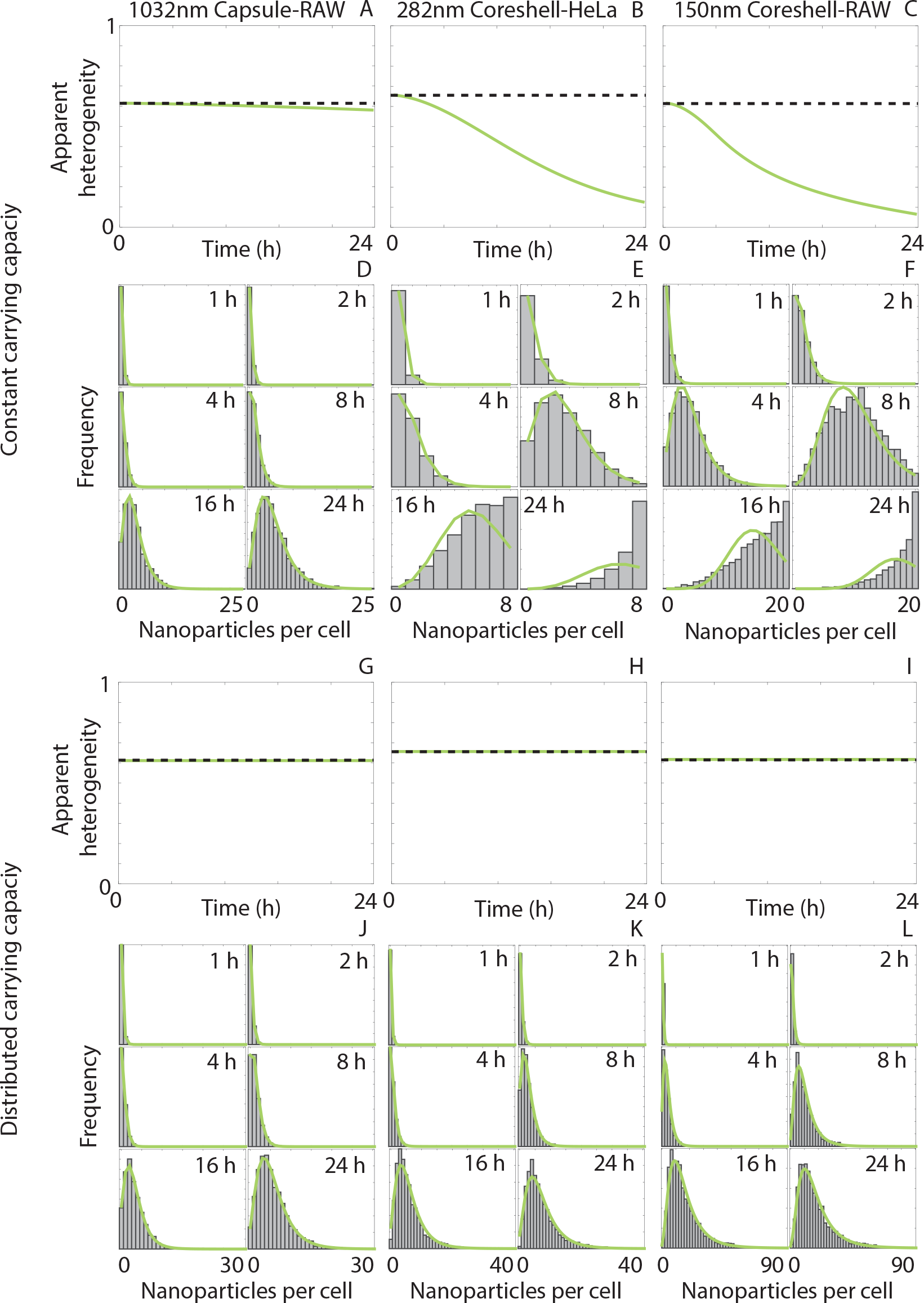
Cell size distribution does not explain changes in heterogeneity. (**A-C**, **G-I**) Evolution of apparent heterogeneity obtained from the modeling framework for three nanoparticle-cell pairs. The affinity parameter for each cell is sampled from a lognormal distribution with a standard deviation corresponding to the dashed black line. The cell carrying capacity parameter is either (**A-C**) constant or (**G-I**) lognormally distributed with a standard deviation corresponding to the dashed black line. (**D-F**, **J-L**) Dosage distributions for the three nanoparticle-cell pairs after 1, 2, 4, 8, 16 and 24 hours obtained from the voxel-based model. The green line corresponds to the Poisson-lognormal distribution that best fits the dosage distribution.

As predicted, the dosage distribution obtained from the model is Poisson-lognormal distributed in the linear association regime, but does not follow the Poisson-lognormal distribution near the carrying capacity. For the simulation of the experiment that remains in the linear association regime (Fig. 4D) the apparent heterogeneity stays close to the “true” heterogeneity in the cell characteristics, that is, the standard deviation of the nanoparticle-cell affinity distribution (Fig. 4A). For the simulation of the two experiments where the number of nanoparticles per cell approaches the carrying capacity (Fig. 4E,F), the apparent heterogeneity decreases as time increases, even though the “true” heterogeneity is constant (Fig. 4B,C).

While this is a potential explanation for the decrease in apparent heterogeneity observed experimentally, the model predicts that the apparent heterogeneity approaches zero as time increases, whereas the apparent heterogeneity in the experimental data appears to approach a finite positive value (Fig. 3). Further, the time dependence in the apparent heterogeneity obtained from the model results from a breakdown in the match between the dosage distribution and the Poisson-lognormal distribution as the number of nanoparticles per cell approaches the carrying capacity (Fig 4D-F). In contrast, the experimental dosage distribution is well-described by the Poisson-lognormal distribution in all cases, even though the number of nanoparticles per cell approaches the carrying capacity in the 282nm coreshell-HeLa and 150nm coreshell-RAW264.7 experiments (Fig. 3). This suggests that the model assumption of a homogeneous carrying capacity is incompatible with the experimental data.

An increase in cell surface area may increase the maximum number of nanoparticles that can associate with a cell. As such, we now impose a lognormal distribution on the cell carrying capacity parameter, where this lognormal distribution has the same standard deviation as the cell surface area distribution. We make the assumption that the nanoparticle-cell affinity and cell carrying capacity parameters are correlated, that is, a cell with higher affinity due to a higher surface area will also have a higher carrying capacity. We perform simulations, again representing the three previously-described experiments, with heterogeneous nanoparticle-cell affinity and heterogeneous cell carrying capacity, and present the results in Fig. 4G-L. The apparent heterogeneity is consistent with the cell surface area heterogeneity in each case (Fig. 4G-I), and the model dosage distributions are all described well by the Poisson-lognormal distribution (Fig. 4J-L). However, the model-predicted apparent heterogeneity due to the variation in cell surface area is constant with time, and therefore insufficient to explain the evolution in the apparent heterogeneity obtained from the experimental data.

### Cell cycle does not introduce time-dependent heterogeneity

The previous results assumed that the cells present in the population are representative of all phases within the cell cycle. To investigate whether the progression through the cell cycle during the experiment affects the apparent heterogeneity, we now incorporate the cell cycle explicitly in our modeling framework. We implement a multistage model of cell cycle progression, where cells exist in one of *m* states, representing different phases of the cell cycle (33). Cells transition between states at rates corresponding to the average time spent in a particular phase. Here we choose *m* = 4, corresponding to the G1, S, G2 and M phases. We consider two approaches for introducing heterogeneity. First, we impose cell heterogeneity via a lognormal distribution as previously. Second, we introduce a phase-specific mean affinity parameter, representing the change in cell size between phases. The standard deviation is independent of phase. When a cell transitions from M phase to G1 phase, and undergoes mitosis, an additional cell is introduced. The daughter cell either inherits the original cell’s affinity parameter, or has the G1 phase affinity parameter, depending on the cell cycle approach considered. The nanoparticle load of the original cell is split evenly between the original cell and the daughter cell.

We perform simulations representing the three experiments described previously, with the addition of the two approaches for modeling heterogeneity via cell cycle progression, ensuring that the mean affinity is consistent with the previous model simulations. The efficient approach used above to obtain the average nanoparticle dose per cell cannot be used here to obtain the apparent heterogeneity as cell behavior must be explicitly described, and as such we revert to the full voxel-based model. For both approaches we observe that the apparent heterogeneity is relatively consistent, though slightly reduced, compared to the true heterogeneity in the cell characteristics (Fig. 5A-C). This slight reduction is associated with mitosis; splitting the nanoparticle load into two cells reduces the number of cells carrying high numbers of nanoparticles. In both cases the apparent heterogeneity is approximately constant over time, unlike the experimental data, which indicates that the cell cycle cannot explain the apparent evolution of heterogeneity in nanoparticle dosage distributions. The choice of modeling approach does not significantly influence either the apparent heterogeneity or the dosage distribution, as highlighted in Fig. 5A-C and Fig. 5D-I, respectively. Therefore, we do not consider the cell cycle in the remainder of this work. The results presented in Fig. 4 and Fig. 5 clearly demonstrate that conclusions drawn from previous investigations (9, 10, 13) into the sources of nanoparticle-cell heterogeneity are incomplete, and are incapable of explaining the observed time-dependence of heterogeneity in nanoparticle dosage.

**Figure 5:**
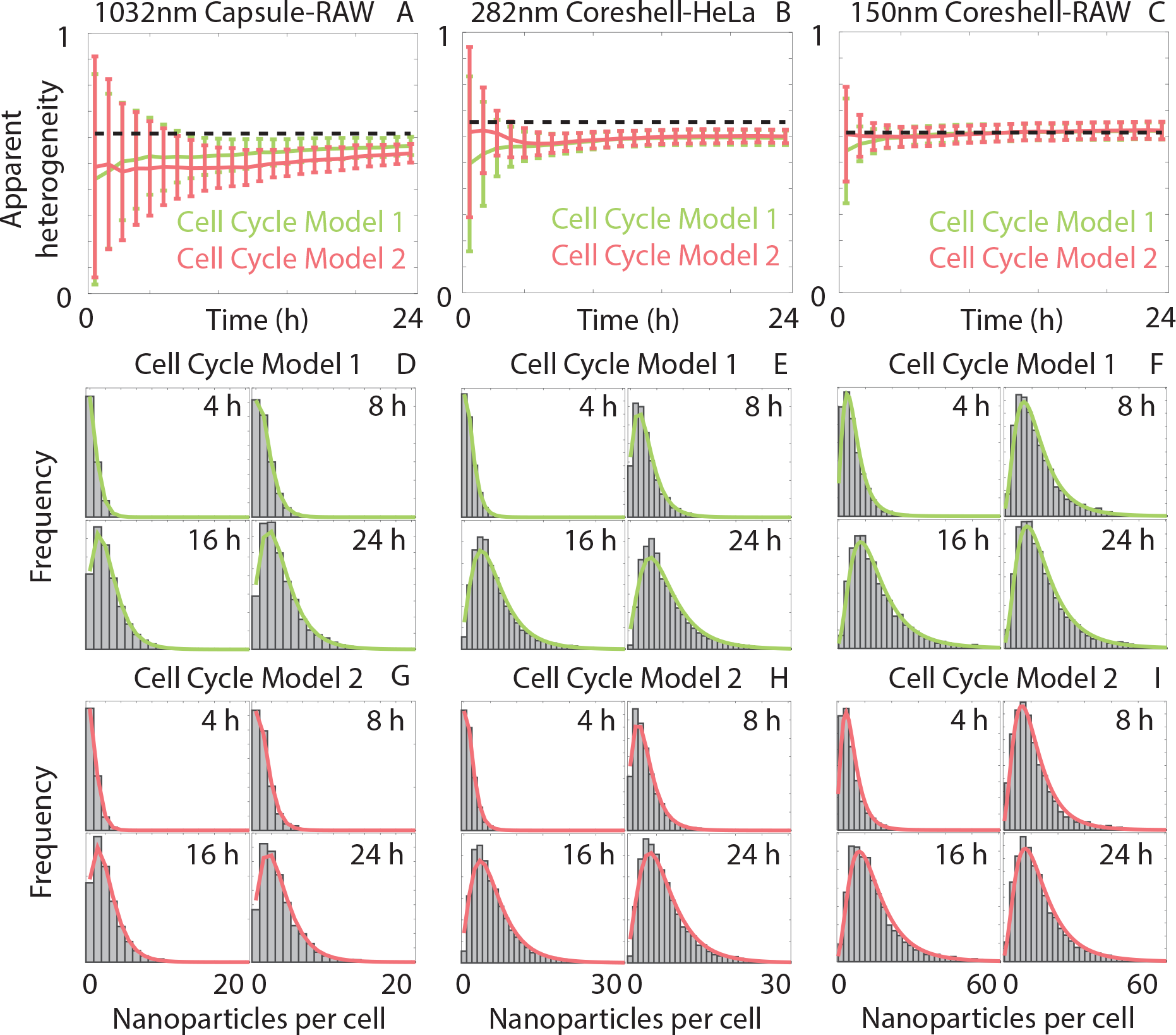
Cell cycle progression does not explain changes in heterogeneity. (**A-C**) Evolution of the average apparent heterogeneity obtained from 200 realizations of the voxel-based framework for the three nanoparticle-cell pairs if the cell cycle is included in the framework via Model 1 (green) and Model 2 (pink). Error bars correspond to one standard deviation. The dashed black line corresponds to the affinity and capacity heterogeneity. (**D-I**) Dosage distributions for the three nanoparticle-cell pairs after 4, 8, 16 and 24 hours obtained from the voxel-based model for (**D-F**) Model 1 and (**G-I**) Model 2. The (**D-F**) green and (**G-I**) pink lines correspond to the Poisson-lognormal distribution that best fit the dosage distribution.

### Interplay between nanoparticle motion, affinity heterogeneity and capacity heterogeneity determines apparent heterogeneity

The association of nanoparticles with cells is eventually restricted by the cell carrying capacity (7). As highlighted by the results in Fig. 4, the heterogeneity in the carrying capacity cannot be neglected. As multiple biological factors may induce heterogeneity in nanoparticle-cell affinity or cell carrying capacity, the amount of heterogeneity may differ between different cell characteristics. Therefore, we now relax the assumption that both the cell carrying capacity and the nanoparticle-cell affinity are distributed with the same degree of heterogeneity. To determine the heterogeneity for both cell carrying capacity and nanoparticle-cell affinity we iteratively calibrate the modeling framework to both the mean nanoparticle dose and the apparent heterogeneity obtained from the experimental data. This allows us to extract estimates of the heterogeneity in the cell carrying capacity and the nanoparticle-cell affinity, while ensuring that the mean dosage is consistent with the experimental data, as shown in Fig. 6. For each experimental dosage curve, we observe that the model accurately describes the mean experimental dose. By varying the nanoparticle-cell affinity heterogeneity and cell carrying capacity heterogeneity independently we are able to match the evolution of the apparent heterogeneity obtained from the experimental data.

**Figure 6:**
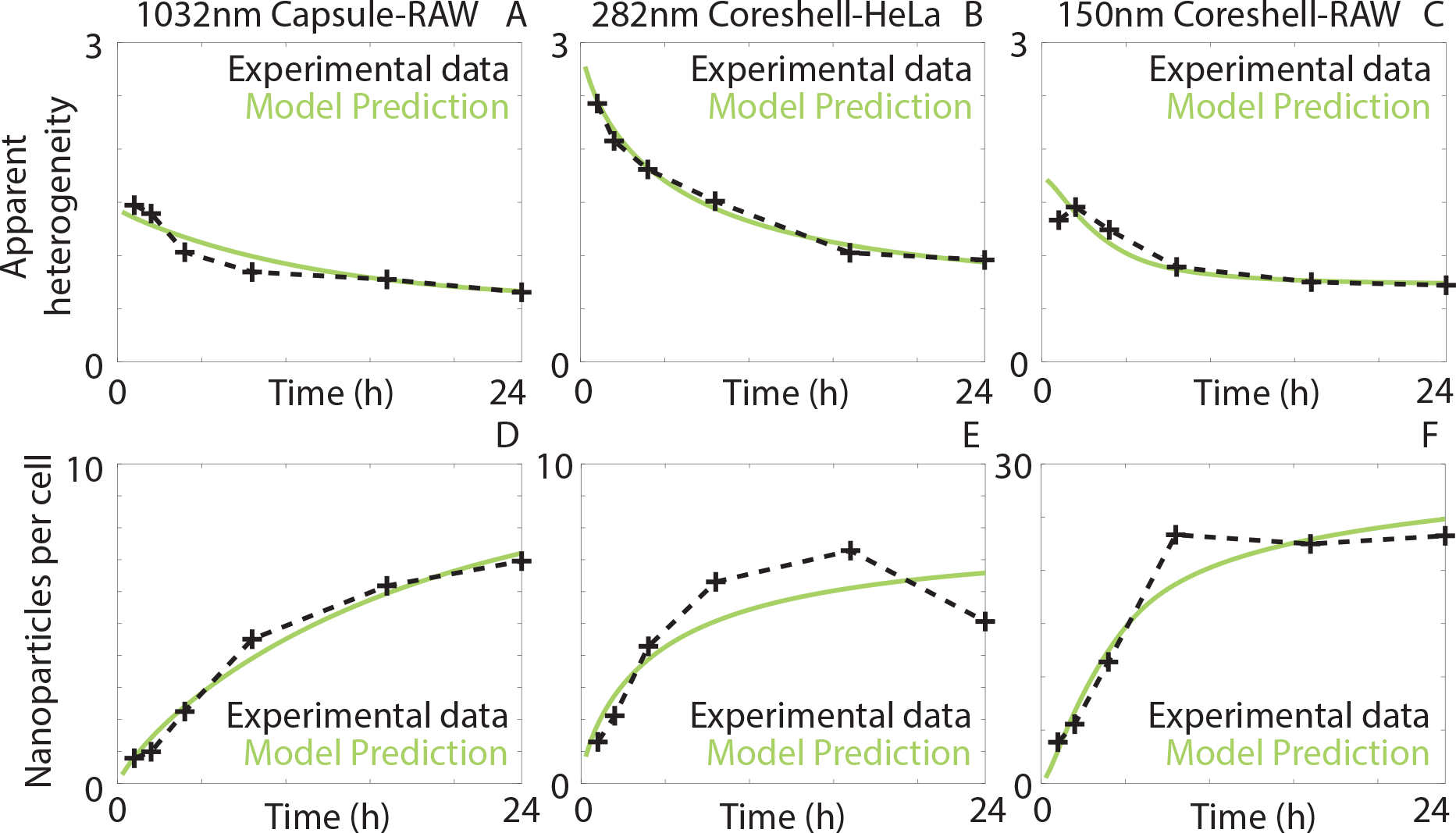
Combining stochastic transport, nanoparticle-cell affinity heterogeneity and cell carrying capacity heterogeneity explains experimentally-observed heterogeneity. Comparison between the model predictions (green) and experimental data (black) for **A-C.** apparent heterogeneity and **D-F.** number of nanoparticles per cell for the three nanoparticle-cell combinations. Here the heterogeneity in nanoparticle-cell affinity is different to the heterogeneity in cell carrying capacity.

At early time, where the rate of nanoparticle association predominantly depends on the nanoparticle-cell affinity, the apparent heterogeneity is close to the “true” heterogeneity in the affinity parameter (1.42, 3.05 and 1.71 in Fig. 6A-C, respectively). As time progresses, the number of associated nanoparticles per cell becomes more dependent on cell carrying capacity. Therefore, we observe that the apparent heterogeneity decreases and approaches the “true” heterogeneity in the cell carrying capacity parameter (0.61, 0.62 and 0.75 in Fig. 6A-C, respectively). This transition demonstrates how these two sources of heterogeneity interact and, consequently, how the “true” heterogeneity manifests itself in the experimental dosage data. All three sources of heterogeneity, namely, the stochastic nanoparticle motion, variation in nanoparticle-cell affinity, and variation in cell carrying capacity, are thus required to explain the heterogeneity in the experimental dosage distributions. Critically, the heterogeneity in the nanoparticle-cell affinity is different to the heterogeneity in the cell carrying capacity. Without this difference in heterogeneity between the two cell parameters, we do not observe the change in the apparent heterogeneity over time.

## Discussion and Conclusions

Experimental data obtained from nanoparticle-cell association assays exhibit variation in the number of nanoparticles associated per cell. We demonstrate that this variation arises from a combination of stochastic nanoparticle transport and heterogeneity in cell characteristics. In particular, heterogeneity in the affinity between nanoparticles and cells, and heterogeneity in the maximum number of associated nanoparticles per cell are shown to be the key biological processes driving variation in experimental data. The amount of variation in the experimental data appears to change with time, but we demonstrate that this is a consequence of different amounts of heterogeneity in these two biological processes.

Uncovering the biological and physical mechanisms that impact the ability of specific cells to associate with, and subsequently internalize, nanoparticles is necessary for informed nanoparticle design (1, 2, 7). The ability to reliably deliver nanoparticles to a target cell population such that the nanoparticles are rapidly internalized has the potential to transform disease treatment and diagnosis (34–36). However, the journey between nanoparticle creation and cellular internalisation is convoluted, and involves a multitude of intertwined biological, physical and chemical processes (1–4, 7). The combination of this complex tapestry with heterogeneous experimental data has thus far inhibited understanding of the key mechanisms governing nanoparticle-cell interactions.

Here we develop a mathematical framework of individual-level nanoparticle-cell interactions that can be used to separate the physical processes governing nanoparticle transport from the biological processes dictating the cellular uptake of nanoparticles. We employ this framework to explain the apparent temporal evolution of heterogeneity within a cell population. This apparent evolution is driven by the interplay between the inherent stochastic motion of the nanoparticles, the heterogeneity in the maximum number of nanoparticles internalized by a particular cell, and the heterogeneity in the affinity between the nanoparticles and the cells. When considered in isolation, these three processes are unable to describe the apparent evolution in heterogeneity present in the experimental data. All three sources of heterogeneity in concert are necessary to explain the experimental data. Further, we demonstrate the impact of the cell cycle is insufficient to explain the heterogeneity in nanoparticle dose. This is in contrast to results obtained from a recent investigation (10), where it was claimed that the combination of stochastic nanoparticle motion and the distribution of cell surface area is sufficient to explain the heterogeneity in nanoparticle dosage. We conclusively demonstrate, using our modeling framework, that this combination does not give rise to heterogeneity that changes over time, as observed experimentally. This has important implications for understanding the impact of heterogeneity, as our modeling approach reveals that the critical biological mechanism inducing heterogeneity in nanoparticle dosage changes over time.

By recognizing that early-time nanoparticle-cell association is driven by nanoparticle-cell affinity and that late-time nanoparticle-cell association is governed by the cell carrying capacity, we provide an intuitive explanation for the apparent evolution of heterogeneity in the experimental data. That is, the apparent heterogeneity is equal to the nanoparticle-cell affinity heterogeneity at early time, and will transition to the cell carrying capacity heterogeneity over time, at a rate corresponding to the nanoparticle transport and association rate.

Extracting reliable estimates of the true heterogeneity in cell characteristics is critical for therapeutic purposes (12, 18, 19). If only a fraction of a cell population appears to internalize a therapeutic dose of nanoparticles, it is crucial to determine whether this is due to stochastic interactions or an underlying biological process. The former implies that increasing the dosage or exposure time will increase the fraction of the population that are effectively treated, whereas the latter suggests that an alternative treatment protocol may be required. Further, the cells that do have lower nanoparticle affinity, and a lower carrying capacity, are more likely to not internalize nanoparticles (Supplementary Information, Fig. S9). This imposes a selective pressure on the cell population toward lower nanoparticle-cell affinity and cell carrying capacity values, as such cells do not respond to the nanoparticle treatment.

The framework presented here demonstrates that mechanistic modeling approaches can be employed to isolate the sources of heterogeneity present in a cell population from nanoparticle-cell association data. Subsequently, this provides insight into whether cell subpopulations exist, and the degree to which such subpopulations are resistant to treatment. More generally, this work highlights how mechanistic insight can be utilized to explain the presence and origin of heterogeneity in experimental data. The influence of cell heterogeneity is a current question interest across a diverse range of fields, including cancer biology (18, 19), molecular biology (37), nanomedicine (10, 13) and microbiology (38). Mechanistic models that incorporate heterogeneity and examine its associated impact, such as the one presented here, may provide the key required to understand the influence of cell heterogeneity.

## Materials and Methods

### Voxel-based Model

While the standard dosage model (Supplementary Information) is effective at calculating the delivered dose for nanoparticle-cell systems, it does not provide information about variation in the nanoparticle dose on a cell-by-cell case. That is, the standard dosage model provides the average number of nanoparticles associated with a single cell, but does not provide a measure of spread. We therefore propose the use of a mathematical model that describes the transport of individual nanoparticles. The combination of diffusion and sedimentation can be represented as a biased random walk, where a nanoparticle undergoes random motion, albeit with a preferential direction corresponding to the direction of sedimentation. However, due to the size discrepancy between nanoparticles and the experimental domain, directly simulating nanoparticle trajectories is extremely computationally demanding. To address this, we consider a voxel-based approach, where the experimental domain is split into discrete subdomains known as voxels. Voxel-based approaches are widely used to model chemical reaction kinetics and cell behavior.

Instead of directly simulating the trajectory of each nanoparticle, the voxel-based model only requires that we simulate the events where nanoparticles transition between voxels. This simplification involves an inherent assumption that nanoparticles are, on average, uniformly distributed throughout each voxel. For a three-dimensional voxel of length *h*, the transition rates from voxel (*i, j, k*), where *i* ∈ [1, *I*], *j* ∈ [1, *J*], *k* ∈ [1, *K*], to a neighbouring voxel for a biased random walk corresponding to sedimentation and diffusion are (21)

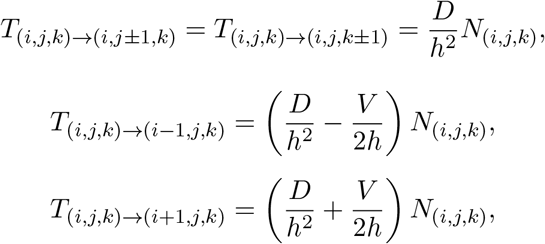

where *N*(_*i,j,k*_) is the number of nanoparticles in voxel (*i, j, k*) and sedimentation occurs in the positive *i* direction. Intuitively, the transition rate is highest in the direction of sedimentation, and lowest in the opposite direction. To account for the zero nanoparticle transport boundary condition at the media-air interface, the transition rate in the negative *i* direction is zero at *i* = 1. That is,

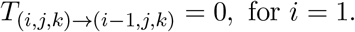

To describe the cell carrying capacity boundary condition at the media-cell interface, we consider the bottom surface of certain voxels at *i* = *I* to be an individual cell. We implement the indicator function *M*_*j,k*_ to denote whether the boundary at (*I, j, k*) contains a cell, that is, *M*_*j,k*_ = 1 if a cell is located at (*I, j, k*), and zero otherwise. The proportion of voxels that contain a cell is equivalent to the level of confluency in the corresponding experiment. We are free to select *h* and a natural choice is to choose *h* such that cross-sectional area of the voxel is equivalent to the surface area of a cell. Further, we are able to track the number of nanoparticles that pass through the media-cell interface at each cell location, which provides detailed information about how the inherently stochastic motion of the nanoparticles influences the spread of the nanoparticle per cell dose. The transition rate corresponding to the cell carrying capacity boundary condition is

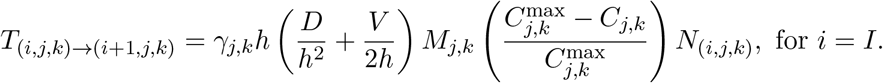

where *γ*_*j,k*_ represents the nanoparticle-cell affinity of the cell at (*I, j, k*) and *C*_*j,k*_ is the number of nanoparticles associated with the cell at (*I, j, k*). The nanoparticle-cell affinity parameter *γ* in the voxel-based model relates to the affinity parameter in the standard model by *γ* = α/D, provided that *D/h* ≫ *V* (Supplementary Information). The remaining boundaries are treated as periodic, that is, when a nanoparticle passes through at *j* = 1, *j* = *J*, *k* = 1, and *k* = *K*, that nanoparticle is removed from the system and another particle is introduced at *j* = *J*, *j* = 1, *k* = *K*, and *k* = 1, respectively. This boundary condition implies that the simulation domain is a representative portion of the experimental domain, and that nanoparticles arrive and leave the simulation domain at, on average, the same rate in the *j* and *k* directions.

To perform realizations of the voxel-based model, we implement an adaptive timestep tau-leap algorithm (39). This is a modified form of Gillespie’s algorithm, implemented to reduce computation time. Initially, we uniformly distribute the nanoparticles throughout the simulation domain. During a timestep of duration *τ*, the number of transition events in each direction for each voxel is sampled from a Poisson distribution according to the appropriate transition rate. As we require that the number of nanoparticles in each voxel is non-negative, if the sampled number of transition events would result in a negative amount of nanoparticles in a voxel, we halve the timestep and sample the number of transition events in the reduced timestep.

### Cell Cycle Model

To describe the cell cycle, we implement a multi-stage model of cell cycle progression, as presented previously (33). At each point in time a cell belongs to one of *m* possible phases of the cell cycle. For the work presented here, we choose *m* = 4, corresponding to the G1, S, G2 and M phases of the cell cycle. Cells undergo stochastic transitions between cell cycle phases via the aforementioned tau-leap algorithm. The possible transitions in the model are from G1 phase to S phase, S phase to G2 phase, G2 phase to M phase, and M phase to G1 phase, with the creation of an additional cell that will also be in G1 phase. These transitions occur at rates denoted *r*_G1_, *r*_S_, *r*_G2_ and *r*_M_, respectively. We obtain estimates of the transitions rates based on the average time a cell spends in each phase. We implement heterogeneity due to cell cycle via two different approaches:

- Approach One. Here the nanoparticle-cell affinity and the cell carrying capacity distributions depend on the distribution of cell area and do not change over the course of the cell cycle. However, cell mitosis still occurs and reduces the nanoparticle load in an individual cell as it produces a daughter cell. After mitosis occurs, the original cell and the daughter cell will have an equal nanoparticle load, which is half of the nanoparticle load before mitosis occurred.
- Approach Two. Here the nanoparticle-cell affinity distribution has a mean value that depends on the cell cycle phase. The standard deviation of the distribution is independent of cell cycle phase. As a cell progresses through the cell cycle, the associated affinity parameter changes. Mitosis occurs as described previously, with the addition that the daughter cell inherits the affinity and cell carrying capacity parameter of the original cell.

### Hybrid Model

While the voxel-based model provides rich detail regarding the location of individual nanoparticles and the nanoparticle load of individual cells, it is computationally intensive to perform realizations of the model. To obtain estimates of the apparent heterogeneity arising from the model, we require a sufficiently large number of cells such that the apparent heterogeneity does not exhibit significant fluctuations due to the stochastic nature of the voxel-based model. While this is feasible, there are more efficient methods, providing that the model is performed in parameter regimes that satisfy several assumptions. We propose a hybrid model, which combines the efficiency of the PDE-based traditional dosage model with the ability of the voxel-based model to provide the nanoparticle dose for individual cells.

The key assumption that the hybrid model relies on is that the location of a cell (relative to other cells) does not affect the ability of that cell to associate with nanoparticles or, equivalently, there is minimal competition between cells for nanoparticles. Mathematically, this is the same as the assumption that the rate of nanoparticle-cell association is small compared to the rate of nanoparticle diffusion out of the voxel. In this case, an individual nanoparticle would likely transition through many boundary voxels before binding to a cell. Under this assumption, the net association rate is simply the sum of all of the individual association rates.

Given that a nanoparticle associates with *any* cell, the probability that it associates with a particular cell located at (*j, k*) is the ratio of the net affinity with that cell to the total affinity of all cells. This can represented as

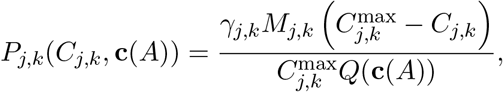

where

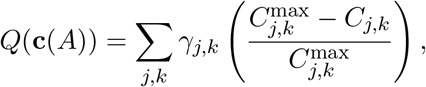

and **c**(*A*) is the set of the number of nanoparticles associated with each cell for a total number of associated nanoparticles *A* = Σ_*j,k*_ C_*j,k*_. That is, (*C*_*j,k*_ ∈ c(*A*) | *j* ∈ (1, *J*), *k* ∈ (1, *K*) s.t. *M*_*j,k*_ = 1). This probability can also be considered the proportion of a single nanoparticle that would associate with a particular cell, given that we know that a nanoparticle has associated with any cell. We note that both *P*_*j,k*_ and *Q* are dependent on the number of nanoparticles associated with each cell, which is, in turn, dependent on the total number of associated nanoparticles. Initially, we know that each cell is associated with zero nanoparticles. The rate of association for each cell, with respect to the total number of associated nanoparticles, can be expressed as a system of ordinary different equations (ODEs)

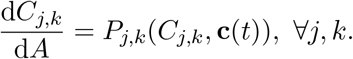

We can solve these equations in terms of the total number of associated nanoparticles; however, this does not provide the evolution of the number of nanoparticles associated per cell in terms of time. To achieve this, we combine this system of ODEs with the traditional dosage model. For a given number of associated nanoparticles *A* = *C*(*t*), we have

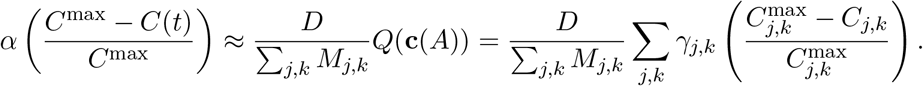

The above simply states that the nanoparticle flux through the boundary due to nanoparticle-cell association is approximately equal to the average of the flux through the boundary due to nanoparticle-cell association for all cells in the system.

The hybrid model is therefore

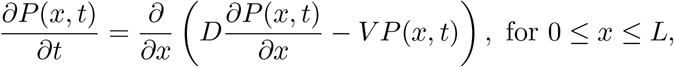

with boundary conditions

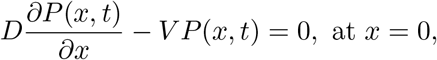

and

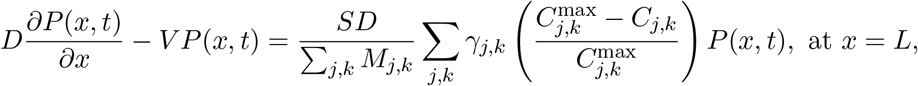

where

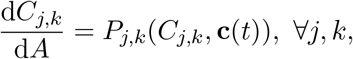

and

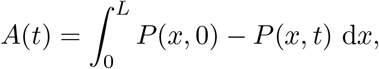

with initial conditions

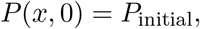

and

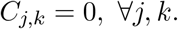

To obtain numerical solutions to the hybrid model we solve the PDE using central spatial differences, the backwards Euler method and Thomas’ algorithm (40). However, to solve the system of ODEs we require a different approach. During an Euler time step of duration Δ*t* the number of nanoparticles that became associated with nanoparticles is Δ*A* = *A*(*t* + Δ*t*) − *A*(*t*). Provided Δ*A* is small, *A*(*t*) provides a close approximation of the number of associated nanoparticles during that time step. After we calculate Δ*A*, we update the system of ODEs via the forward Euler method. The method to solve the system is therefore: (i) solve the PDE from *t* to Δ*t*; (ii) calculate Δ*A* from the new solution values; (iii) solve the system of ODEs from *A*(*t*) to *A*(*t* + Δ*t*); and (iv) repeat.

### Poisson-lognormal Distribution

Nanoparticles arrive at the cell-media interface at a rate proportional to the transport properties of the nanoparticles. Away from the cell carrying capacity, the binding to the cell surface and subsequent internalization occurs at a rate directly proportional to the affinity parameter *α*. This binding is probabilistic and described by the Poisson distribution; the probability that *k* nanoparticles are associated with a cell after a given time is

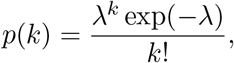

where *λ* is the average number of nanoparticles associated with a cell after that time. As the number of associated nanoparticles is directly proportional to the affinity, λ ∝ α (as per the flux boundary condition). The nanoparticle dosage for the cell population is *N*_cell_ samples from *p*(*k*), where *N*_cell_ is the number of cells in the population.

Now consider the heterogeneous case where *α* is lognormally distributed. That is, each cell has an affinity parameter, *α*_*i*_, that is sampled from the lognormal distribution with mean value 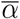 and standard deviation *σ*_*α*_. For an individual cell, the probability mass function that *k* nanoparticles, given an affinity parameter *α*_*i*_, are associated with that cell is

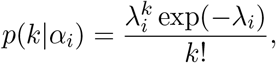

where *λ*_i_ ∝ α_i_. The probability distribution function for *k* nanoparticles associated with a particular cell with affinity parameter *α_i_* is

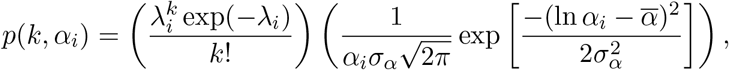

The dosage distribution for a representative population of cells is therefore

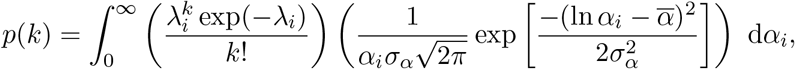

which we refer to as the Poisson-lognormal distribution (29).

### Experimental Approach

HeLa and RAW 264.7 cells were seeded in standard 12-well culture plates at a density of 10^5^ cells per well. Cells were incubated in 1 mL total of media, including 10% fetal bovine serum and fluorescently-labeled particles for the requisite experiments. Introduced particle concentration was 100:1 particles per cell for all time points. After incubation, cells were washed gently with sterile phosphate buffered saline to remove un-associated particles, then detached from wells through the administration of trypsin. Trypsin was deactivated through the administration of growth media, cells were then re-suspended in phosphate buffered saline. Cells (now in suspension) were then analyzed through flow cytometry to determine number of particles associated (bound or internalized) to each cell analyzed. A minimum of 10^4^ cells were analyzed for each sample. Time course incubation data up to 24 hours was obtained with independent samples for each time point, which were analyzed at roughly the same time. Time point “0” (i.e., incubated with particles for 0 minutes) was obtained by analyzing cells that had not been incubated with particles (i.e., a blank control). As per (7), the number of associated particles for cell *i* at time *t*, *N*_*i*_(*t*), is obtained via

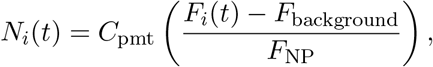

where *C*_pmt_ is a correction factor that accounts for photomultiplier voltage settings (details in (7)), *F*_*i*_(*t*) is the fluorescence of cell *i*, which has been incubated with nanoparticles, *F*_background_ is the median fluorescence intensity of cells in the absence of nanoparticles and *F*_NP_ is the median fluorescence intensity of nanoparticles in solution.

Polymer particles were synthesized through layer-by-layer assembly on nanoparticle templates of gold (Au, 150nm particles) or silica (SiO2, 282nm, 1032nm particles). Briefly, a layer of disulfide-stabilized poly(methacrylic acid) (PMASH) was adsorbed onto the surface of particles, followed with a layer of poly(N-vinyl pyrrolidone) (PVPON) which adhere to the PMASH layers through hydrogen bonding. This process was repeated for a total of 4 polymer bilayers. Crosslinking of disulfides within the PMASH layers was then accomplished to stabilize the particles, and PVPON was removed through increasing the pH; leading to PMASH core-shell particles. For capsules, templates were removed using potassium cyanide (Au templates) or hydrofluoric acid (SiO2 templates). Characterization of the nanoparticles indicates monodispersity. Full details of synthesis and experiments are available in (7).

## Data and code availability

Certain experimental data used in this analysis have been previously published (7) and raw data are available at https://figshare.com/projects/Invitrocell-particleassociation/59162. The code used to implement the mathematical framework is available at https://github.com/DrStuartJohnston/nanoparticle-cell-interactions.

## Author contributions

STJ conceived the study. STJ, MF and EJC designed the numerical experiments. STJ designed and performed the analysis. STJ wrote the manuscript. STJ, MF and EJC edited the manuscript. All authors gave final approval for publication.

## Acknowledgements

This research was in part conducted and funded by the Australian Research Council Centre of Excellence in Convergent Bio-Nano Science and Technology (project no. CE140100036).

## Traditional Dosage Model

The standard mathematical approach for describing nanoparticle transport is through the use of a partial differential equation (PDE) [1, 3, 5, 6]. This PDE states that the time evolution of the nanoparticle concentration is due to the gradient of a combination of diffusive flux (random motion) and advective flux (sedimentation). Mathematically, this can be stated as

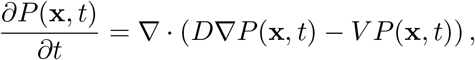

where *P* (**x**, *t*) is the nanoparticle concentration, *D* is the diffusivity and *V* is the sedimentation velocity [5]. The diffusivity of a nanoparticle arises from the Stokes-Einstein equation:

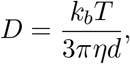

where *k*_*b*_ is the Boltzmann constant, *T* is the temperature of the media, *η* is the dynamic viscosity of the media and *d* is the nanoparticle diameter. The sedimentation velocity of a nanoparticle can be obtained from Stokes’ Law:

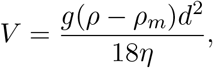

where *g* is the gravitational acceleration constant, *ρ* is the nanoparticle density and *ρ*_m_ is the media density. A standard simplifying assumption for nanoparticle-cell association experiments with static culture media is that the nanoparticle concentration only varies in the vertical dimension, reducing the model to a single spatial dimension [3, 5, 6]:

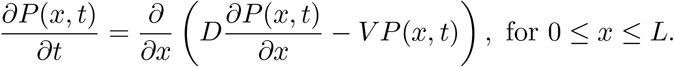

The interface between the cell monolayer and culture media (*x* = *L*) and the interface between the media and the air (*x* = 0) provide natural boundary conditions for the model. Intuitively, there will be no transport of nanoparticles from the media into the air, which can be represented mathematically as

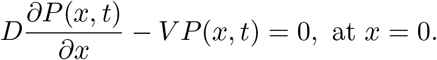

The boundary condition at the media-cell interface is more complicated. Models have been proposed where the nanoparticle concentration at the interface is zero, which involves the assumption that any nanoparticle arriving at the cell monolayer immediately associates with a cell and is removed from the media [6]. Other models make the assumption that there is a linear flux of nanoparticles into the cell monolayer, governed by a parameter that represents the nanoparticle-cell association rate [3]. More complicated boundary conditions, including representing nanoparticle-cell association as a Langmuir binding event, have also been proposed [1].

More recently, a thorough investigation of multiple nanoparticle-cell systems by Faria *et al.* [3] demonstrated that the media-cell interface is best approximated using a cell carrying capacity model, where cells are able to associate with nanoparticles up to a threshold nanoparticle number. Mathematically, this can be represented as

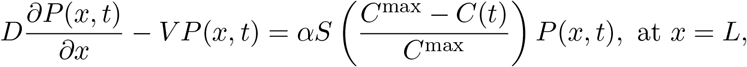

where *α* (m/s) represents the affinity between nanoparticles and cells, *S* is the portion of the culture dish covered by the cell monolayer (level of confluence), *C*^max^ is the nanoparticle carrying capacity of the cells and *C*(*t*) is the number of nanoparticles associated with cells. We obtain this via

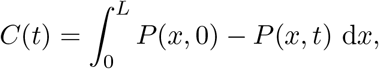

which provides the difference between the number of nanoparticles initially in the system, and the number of nanoparticles in the system at time *t*.

The number of nanoparticles in the system is obtained from the initial condition *P* (*x*, 0) = *P*_0_. That is, there is a specified nanoparticle concentration that is independent of location, consistent with a well-mixed nanoparticle suspension.

To solve the traditional dosage model we apply finite difference techniques, using central difference approximations for the spatial derivatives with node spacing Δ*x* [7]. We implement the backwards Euler method for the temporal derivative, and solve the resulting tridiagonal system of algebraic equations using Thomas’ algorithm [7].

## Results

### Model Equivalence

Consider the transition probabilities for a nanoparticle located at (*x*(*t*), *y*(*t*), *z*(*t*)) = (*ih, jh, kh*).

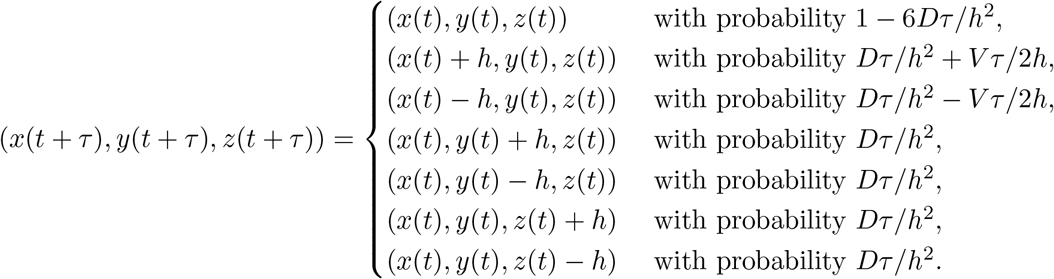

For a voxel on the cell-media boundary, at *x* = *Ih*, assume the nanoparticle is removed from the system (associated with a cell) with probability *P*_assoc_*h* if it attempts to transition through the cell-media boundary. The transition probabilities are therefore

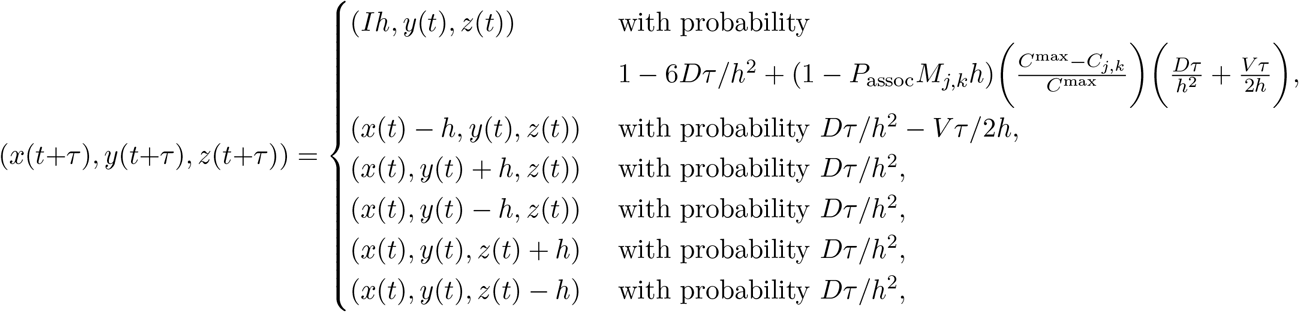

and removed from the system with probability *P*_assoc_*M*_*j,k*_*h*(*Dτ*/*h*^2^ + *Vτ*/2*h*)(*C*^max^ − *C*_*j,k*_)/*C*^max^.

Consider now the evolution of the number of nanoparticles in the voxel (*i, j, k*)

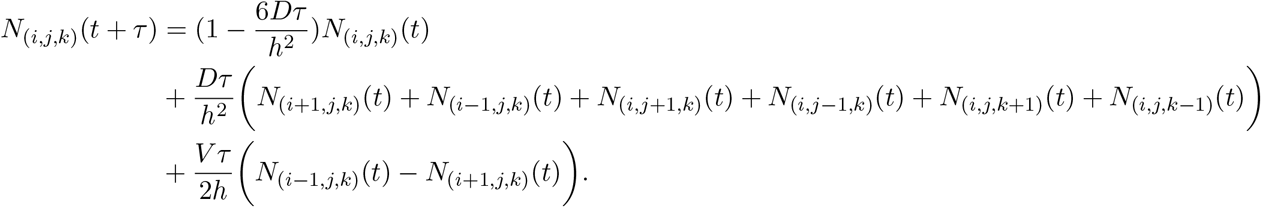

Rearranging, we obtain

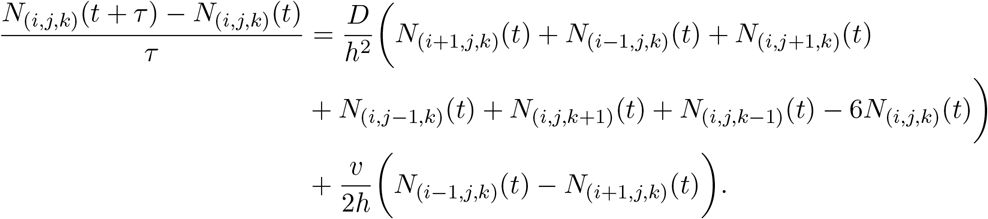

Taking the limit *τ* → 0, *h* → 0 [2], we recover the traditional dosage model.

For the cell-media boundary, the evolution of the number of nanoparticles in a boundary voxel is

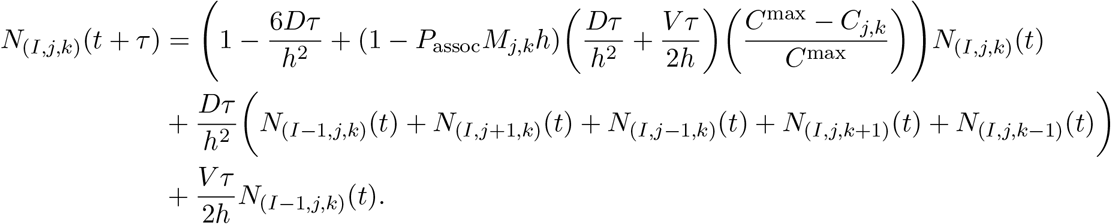

Upon rearranging, we obtain

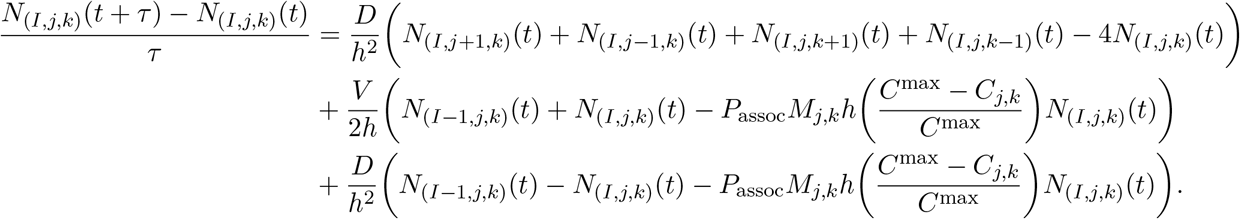

This is equivalent to

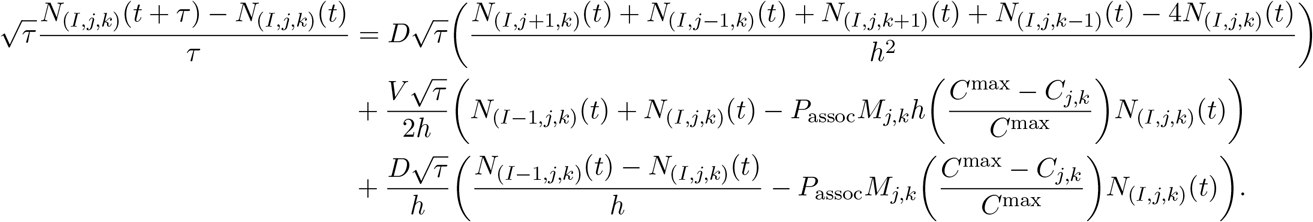

Taking the limit *τ* → 0, *h* → 0 such that 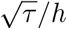 is finite [2], we obtain

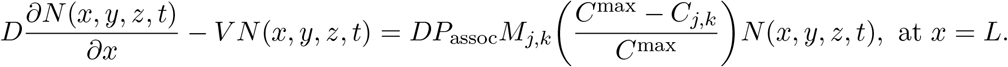

This is consistent with the boundary condition in the traditional dosage model provided

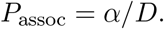

**Table 1:**
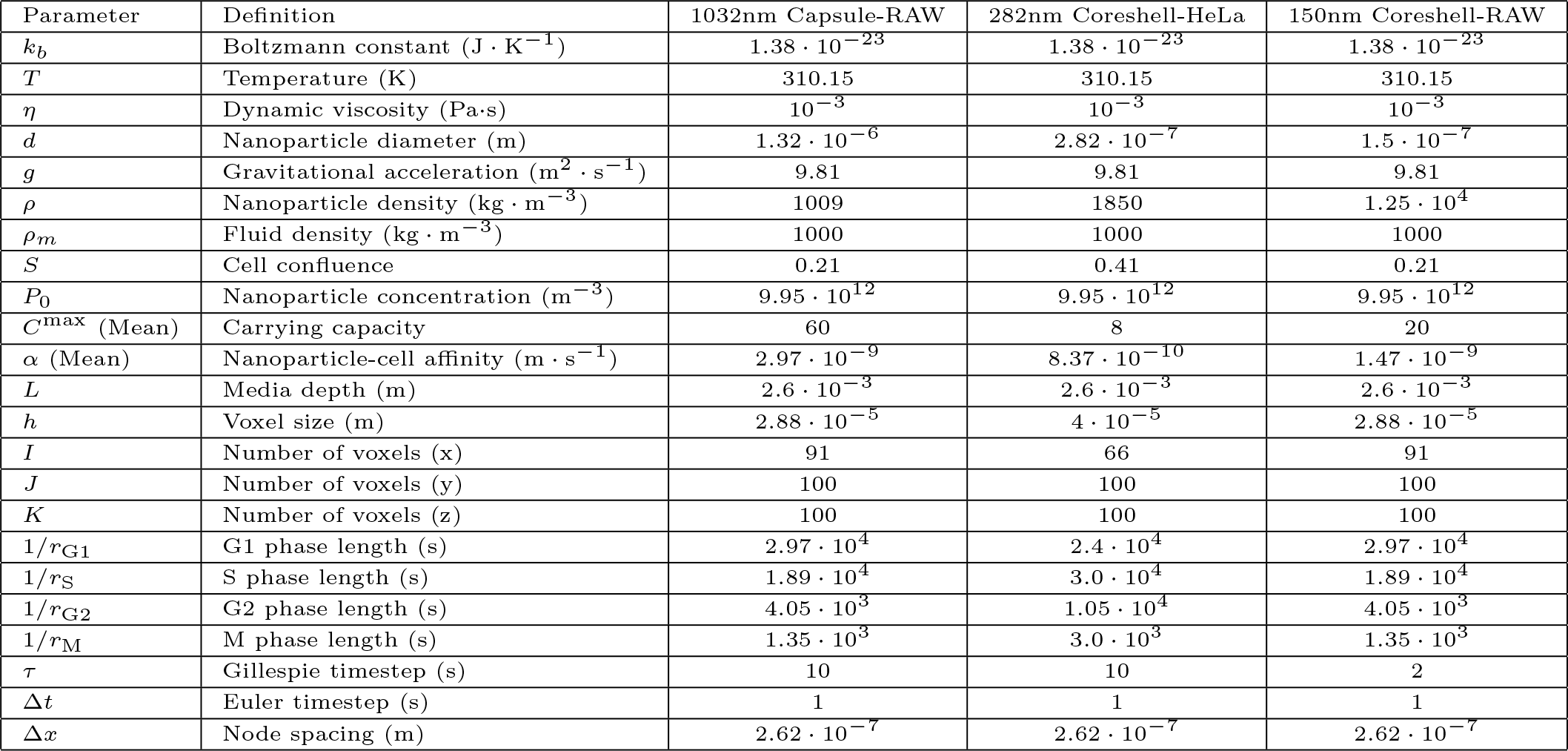
Table of parameters for both the hybrid dosage model and voxel-based model. Definitions of individual parameters are as per the text. Parameters *T*, *η*, *d*, *ρ*, *ρ*_*m*_, *S*, *P*_0_, *C*_max_, *α*, *L* are obtained from [3]. Parameters *r*_G1_, *r*_S_, *r*_G2_, *r*_M_ are estimated from [4] for the 282nm Coreshell-HeLa experiments and from [8] for the 1032nm Capsule-RAW and 150nm Coreshell-RAW experiments. All other parameters are well-established or are numerical solver (user-selected) parameters.

### Model Parameters

The parameters used to obtain numerical solutions to the hybrid dosage model and the voxel-based model are presented in Table 1 for each experiment. Note that for These parameters are used in all figures in the main document, with the exception of Figure 7, where the nanoparticle-cell affinity and cell carrying capacity are obtained via nonlinear least squares. For Figure 7, the nanoparticle-cell affinity parameters are *α* = 3.79.10^−8^, *α* = 3.77.10^−7^, and *α* = 1.48·10^−8^ for the 1032nm Capsule-RAW, 282nm Coreshell-Hela and 150nm Coreshell-RAW experiments, respectively. The cell carrying capacity parameters are *C*^max^ = 9.59, *C*^max^ = 7.69, and *C*^max^ = 28.42 for the 1032nm Capsule-RAW, 282nm Coreshell-Hela and 150nm Coreshell-RAW experiments, respectively.

### Additional Results

In Figure S1, we show that the dose obtained from the model is only Poisson distributed in the linear association regime (Fig. S1A-D). If the number of nanoparticles per cell approaches the carrying capacity, the predicted dosage distribution is not well described by the Poisson distribution (Fig. S1E-F). Comparing the dosage distribution obtained from the model (Fig. S1A-F) with the experimental data (Fig. S1G-L), we observe that the experimental data is overdisperse compared to the model predictions.

In Figure S2 we demonstrate that the number of nanoparticles associated with cells is consistent with the results obtained from traditional dosage model.

In Figure S3 we demonstrate that the apparent heterogeneity obtained from the hybrid model matches the apparent heterogeneity obtained from the voxel-based model.

In Figures S4-S8, we present the difference between the Poisson distribution (Figure S4) or Poisson-lognormal distribution (Figure S5-S8) and the dosage distribution obtained from the hybrid model. The difference is defined as

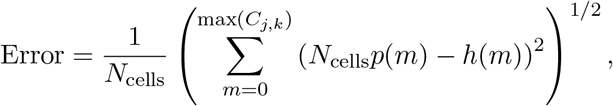

where *N*_cells_ is the number of cells in the simulation, *p*(*m*) is the probability density function for the appropriate distribution and *h*(*m*) is the number of cells that are associated with *m* nanoparticles. Error values that grow with time indicate that the assumptions regarding the distribution are inappropriate. As expected, the error in Figure S4 and Figure S6 grows for the 282nm Capsule-HeLa and 150nm Capsule-RAW experiments, due to the constant carrying capacity. For all other experiments we observe that error is consistently low and hence the assumption of the Poisson-lognormal distribution appears appropriate.

**Figure S1:**
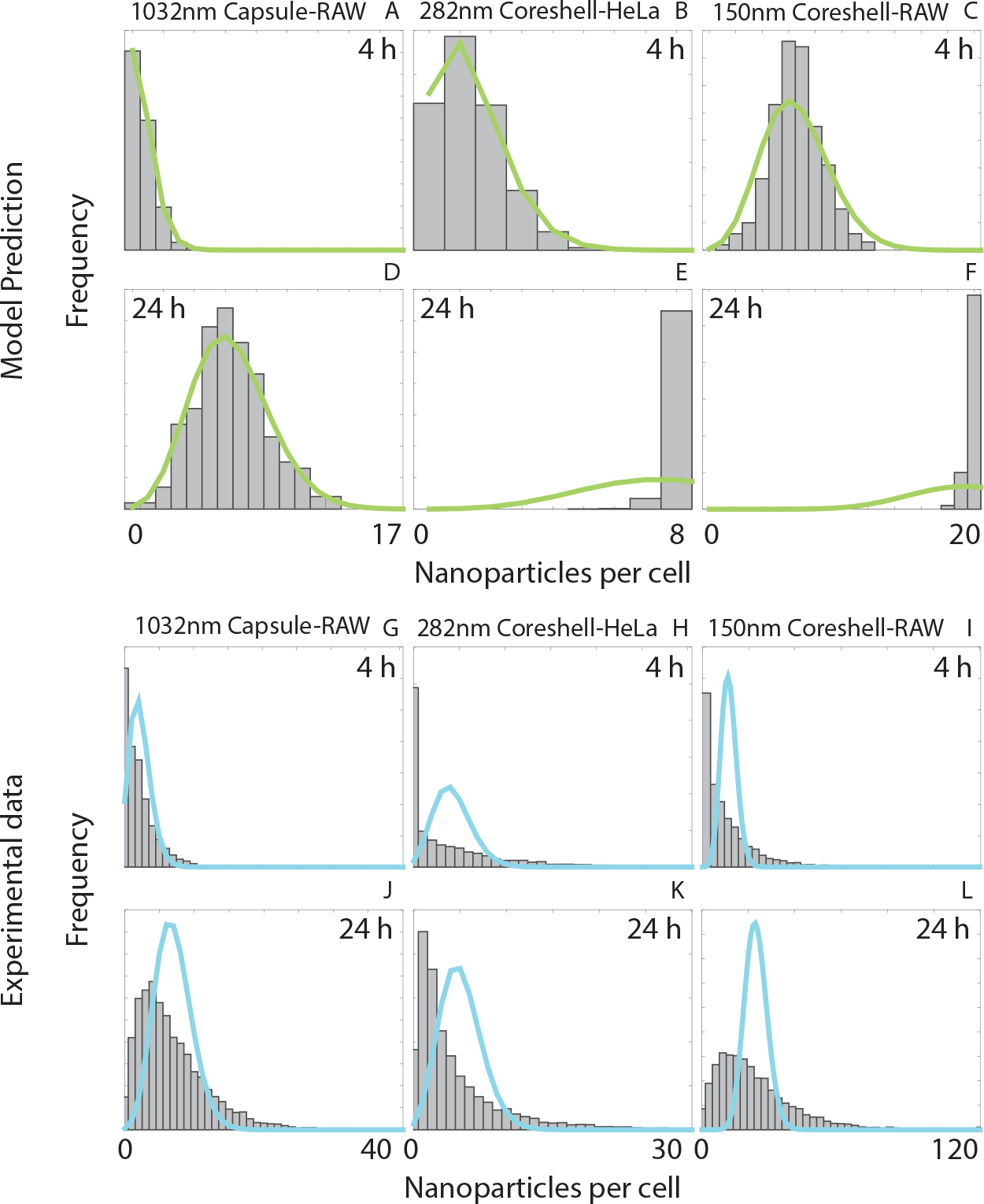
Stochastic motion does not account for all variation in dosage distributions. Histograms of the number of associated nanoparticles per cell for three different nanoparticle-cell pairs after (**A-C**) 4 hours and (**D-F**) 24 hours obtained from the voxel-based modelling framework, and after (**G-I**) 4 hours and **J-L.** 24 hours from the experimental data. The (**A-F**) green and (**G-L**) cyan lines correspond to the Poisson distribution that best fits the dosage distribution.

**Figure S2:**
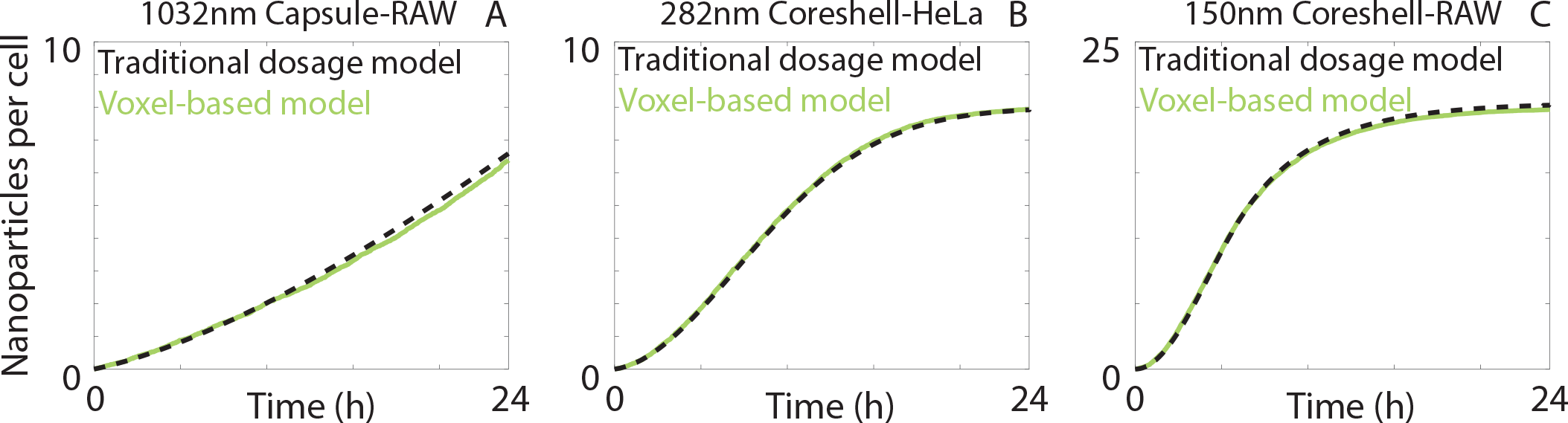
Comparison in mean nanoparticle dose between the traditional dosage model and the voxel-based model.

**Figure S3:**
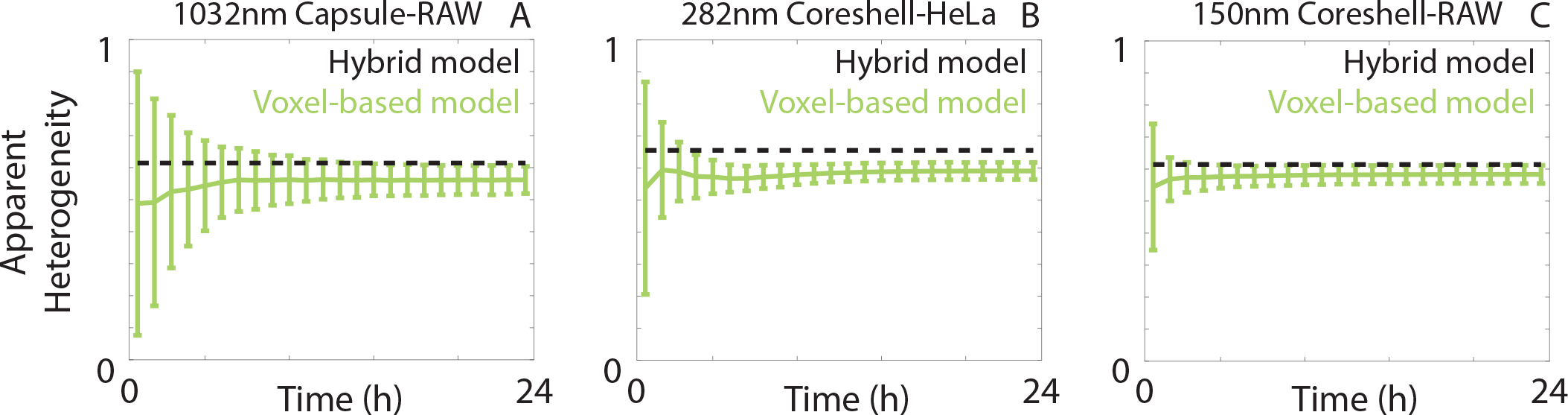
Comparison in apparent heterogeneity between the hybrid model and two hundred realisations of the voxel-based model. Error bars correspond to one standard deviation.

**Figure S4:**
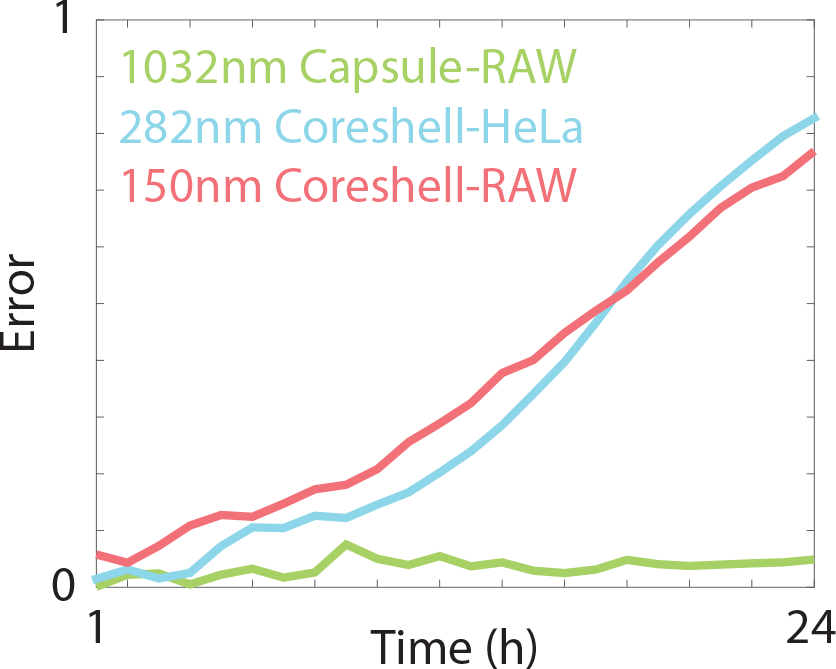
Difference between the Poisson distribution and the model dosage distribution in Figure S1.

**Figure S5:**
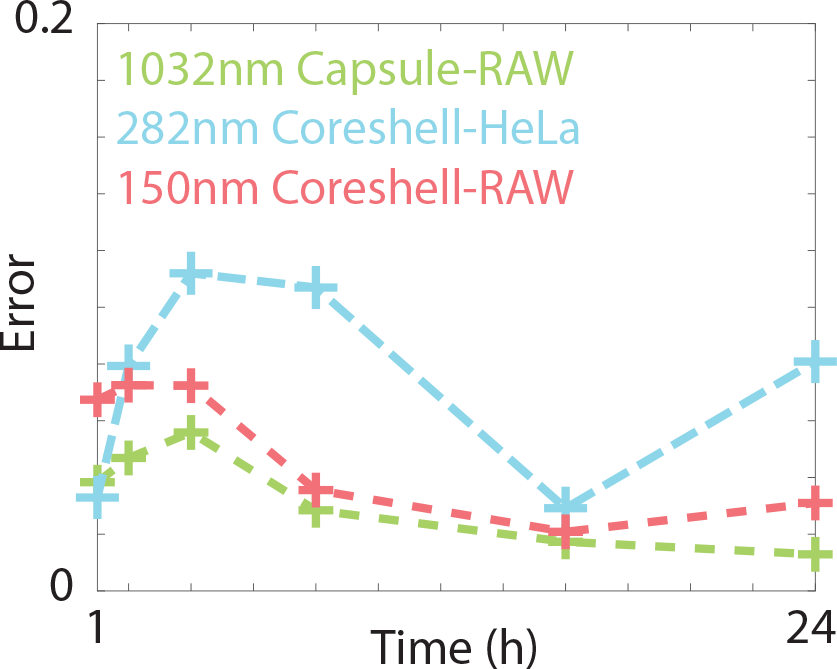
Difference between the Poisson-lognormal distribution and the experimental dosage distribution in Figure 3 (main document).

**Figure S6:**
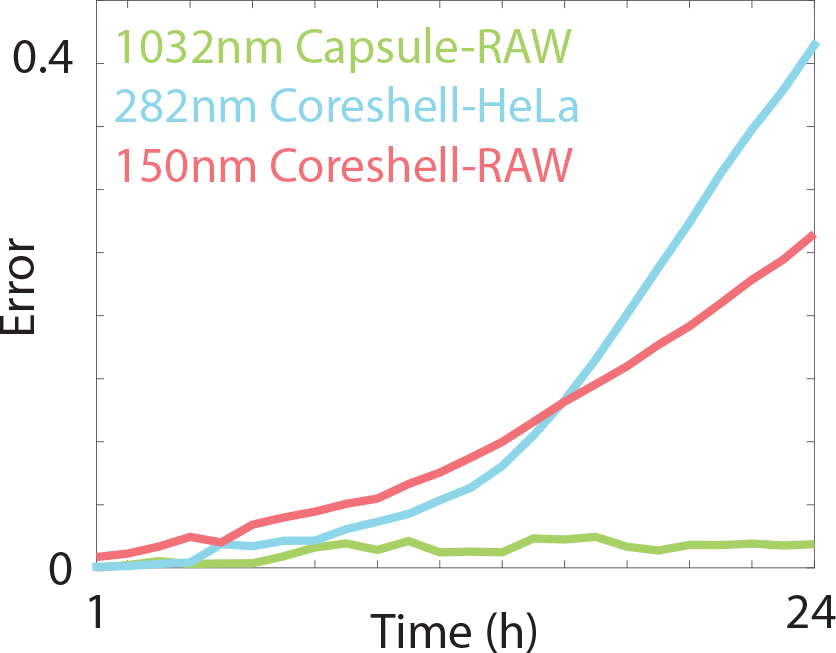
Difference between the Poisson-lognormal distribution and the model dosage distribution in Figures 4D-F (main document).

**Figure S7:**
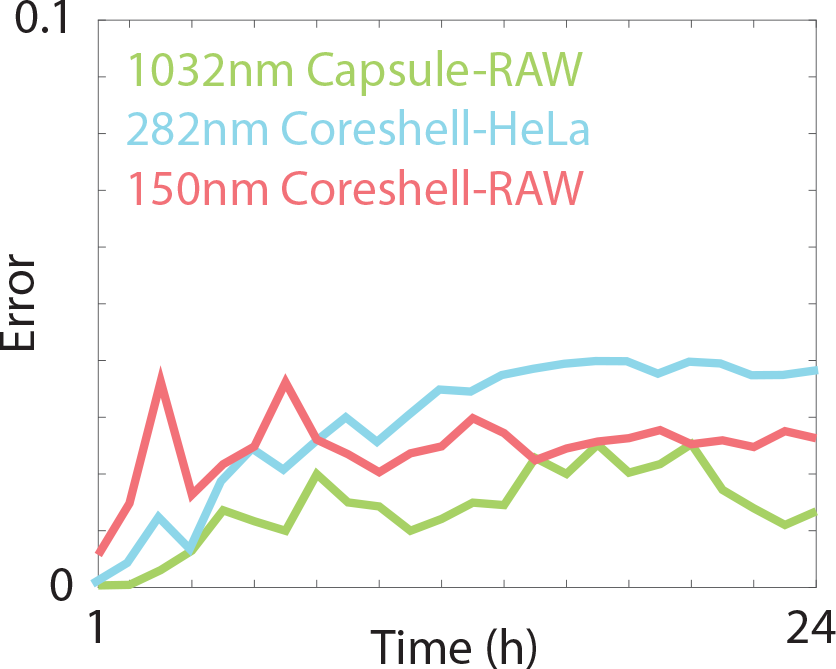
Difference between the Poisson-lognormal distribution and the model dosage distribution in Figures 4J-L (main document).

**Figure S8:**
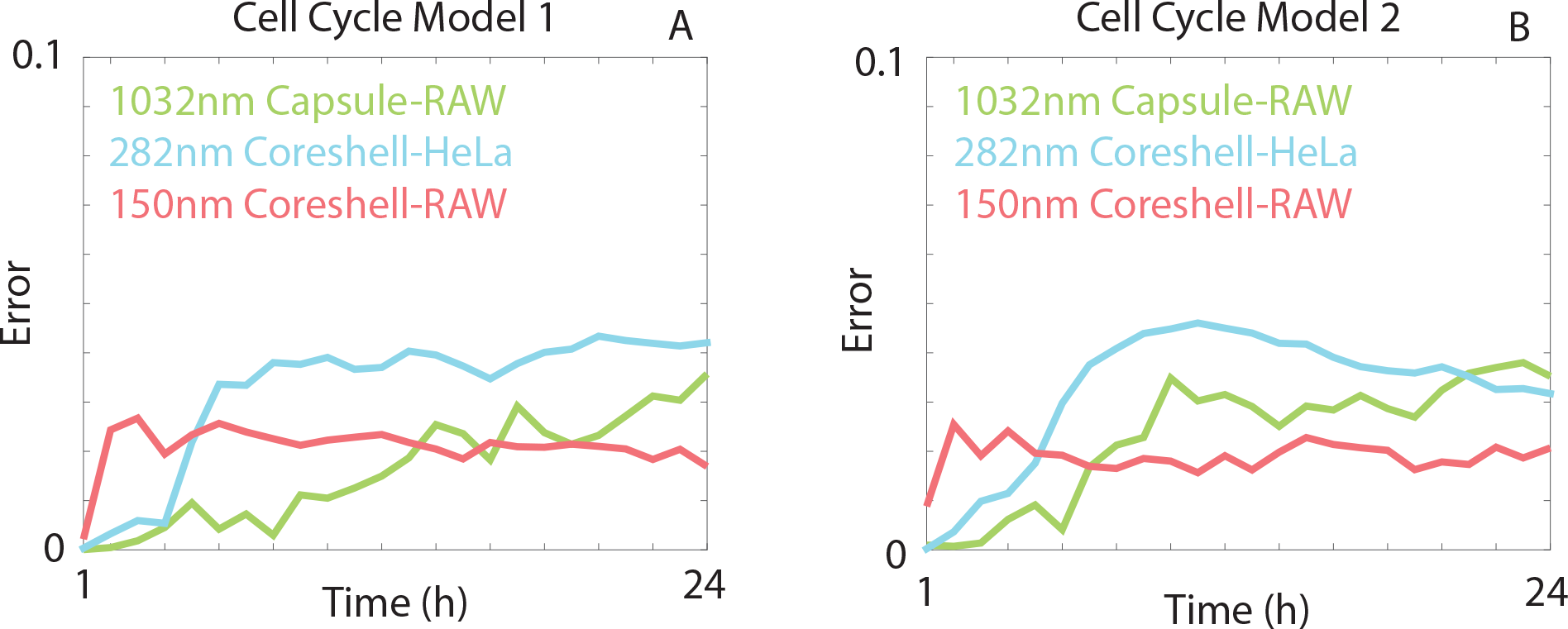
Difference between the Poisson-lognormal distribution and the model dosage distribution in Figure 5 (main document) for (A) Cell Cycle Model 1 and (B) Cell Cycle Model 2.

**Figure S9:**
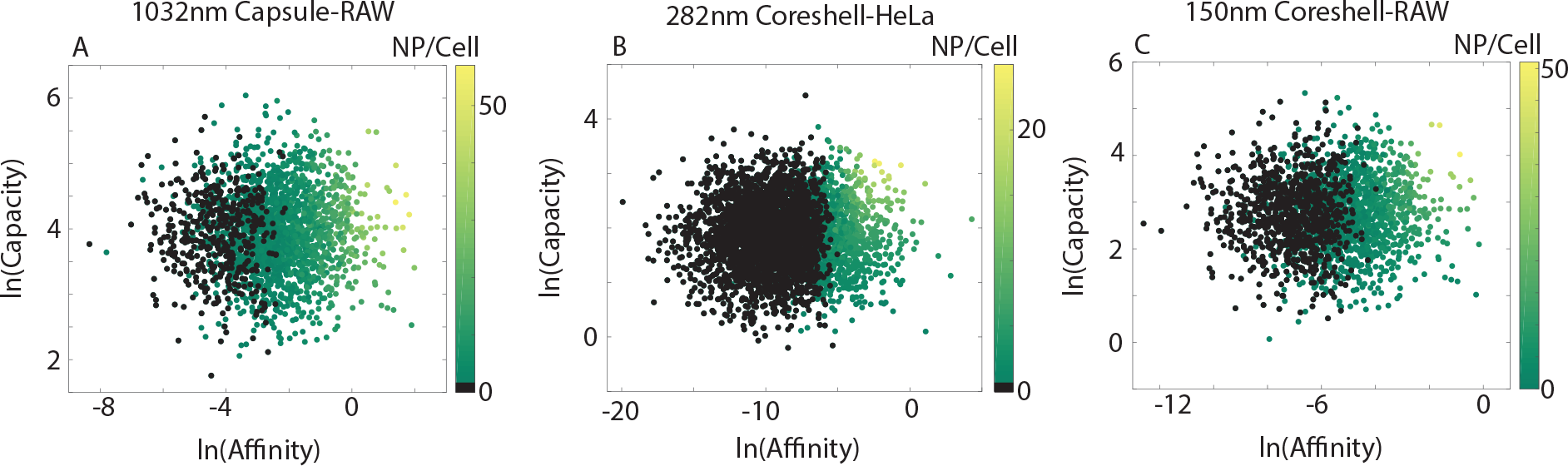
Scatter plots highlighting the relationship between nanoparticle-cell affinity, cell carrying capacity and the number of associated nanoparticles. Black circles correspond to cells with nanoparticle-cell affinity and cell carrying capacity values that resulted in zero associated nanoparticles.

In Figure S9 we present scatter plots of nanoparticle-cell affinity values and carrying capacity values, as well as the corresponding number of associated nanoparticles per cell at the end of 24 hours, as predicted by the voxel-based model. Parameter values that correspond to cells that do not associate with any nanoparticles are coloured in black. We observe that such cells typically have lower nanoparticle-cell affinity values and low cell carrying capacity values, which would impose a selective pressure towards these values if the nanoparticles contained a therapeutic agent.

