## Supplementary Information for "Isolating the sources of heterogeneity in nanoparticle-cell interactions"

### Isolating the sources of heterogeneity in nanoparticle-cell interactions Supplementary Information

#### Traditional Dosage Model

The standard mathematical approach for describing nanoparticle transport is through the use of a partial differential equation (PDE) [1, 3, 5, 6]. This PDE states that the time evolution of the nanoparticle concentration is due to the gradient of a combination of diffusive flux (random motion) and advective flux (sedimentation). Mathematically, this can be stated as

$$\frac{\partial P(\mathbf{x},t)}{\partial t} = \nabla \cdot \left( D\nabla P(\mathbf{x},t) - VP(\mathbf{x},t) \right),$$

where $P(\mathbf{x},t)$ is the nanoparticle concentration, $D$ is the diffusivity and $V$ is the sedimentation velocity [5]. The diffusivity of a nanoparticle arises from the Stokes-Einstein equation:

$$D = \frac{k_b T}{3\pi\eta d},$$

where $k_b$ is the Boltzmann constant, $T$ is the temperature of the media, $\eta$ is the dynamic viscosity of the media and $d$ is the nanoparticle diameter. The sedimentation velocity of a nanoparticle can be obtained from Stokes' Law:

$$V = \frac{g(\rho - \rho_m)d^2}{18\eta},$$

where $g$ is the gravitational acceleration constant, $\rho$ is the nanoparticle density and $\rho_m$ is the media density. A standard simplifying assumption for nanoparticle-cell association experiments with static culture media is that the nanoparticle concentration only varies in the vertical dimension, reducing the model to a single spatial dimension [3, 5, 6]:

$$\frac{\partial P(x,t)}{\partial t} = \frac{\partial}{\partial x}\left( D\frac{\partial P(x,t)}{\partial x} - VP(x,t) \right), \text{ for } 0 \le x \le L.$$

The interface between the cell monolayer and culture media ($x = L$) and the interface between the media and the air ($x = 0$) provide natural boundary conditions for the model. Intuitively, there will be no transport of nanoparticles from the media into the air, which can be represented mathematically as

$$D\frac{\partial P(x,t)}{\partial x} - VP(x,t) = 0, \text{ at } x = 0.$$

The boundary condition at the media-cell interface is more complicated. Models have been proposed where the nanoparticle concentration at the interface is zero, which involves the assumption that any nanoparticle arriving at the cell monolayer immediately associates with a cell and is removed from the media [6]. Other models make the assumption that there is a linear flux of nanoparticles into the cell monolayer, governed by a parameter that represents the nanoparticle-cell association rate [3]. More complicated boundary conditions, including representing nanoparticle-cell association as a Langmuir binding event, have also been proposed [1].

More recently, a thorough investigation of multiple nanoparticle-cell systems by Faria *et al.* [3] demonstrated that the media-cell interface is best approximated using a cell carrying capacity model, where cells are able to associate with nanoparticles up to a threshold nanoparticle number. Mathematically, this can be represented as

$$D\frac{\partial P(x,t)}{\partial x} - VP(x,t) = \alpha S\left(\frac{C^{\max} - C(t)}{C^{\max}}\right)P(x,t), \text{ at } x = L,$$

where $\alpha$ (m/s) represents the affinity between nanoparticles and cells, $S$ is the portion of the culture dish covered by the cell monolayer (level of confluence), $C^{\max}$ is the nanoparticle carrying capacity of the cells and $C(t)$ is the number of nanoparticles associated with cells. We obtain this via

$$C(t) = \int_0^L P(x,0) - P(x,t)\ \mathrm{d}x,$$

which provides the difference between the number of nanoparticles initially in the system, and the number of nanoparticles in the system at time $t$.

The number of nanoparticles in the system is obtained from the initial condition $P(x,0) = P_0$. That is, there is a specified nanoparticle concentration that is independent of location, consistent with a well-mixed nanoparticle suspension.

To solve the traditional dosage model we apply finite difference techniques, using central difference approximations for the spatial derivatives with node spacing $\Delta x$ [7]. We implement the backwards Euler method for the temporal derivative, and solve the resulting tridiagonal system of algebraic equations using Thomas' algorithm [7].

### Results

#### Model Equivalence

Consider the transition probabilities for a nanoparticle located at $(x(t), y(t), z(t)) = (ih, jh, kh)$.

$$(x(t+\tau), y(t+\tau), z(t+\tau)) = \begin{cases} (x(t), y(t), z(t)) & \text{with probability } 1 - 6D\tau/h^2, \\ (x(t)+h, y(t), z(t)) & \text{with probability } D\tau/h^2 + V\tau/2h, \\ (x(t)-h, y(t), z(t)) & \text{with probability } D\tau/h^2 - V\tau/2h, \\ (x(t), y(t)+h, z(t)) & \text{with probability } D\tau/h^2, \\ (x(t), y(t)-h, z(t)) & \text{with probability } D\tau/h^2, \\ (x(t), y(t), z(t)+h) & \text{with probability } D\tau/h^2, \\ (x(t), y(t), z(t)-h) & \text{with probability } D\tau/h^2. \end{cases}$$

For a voxel on the cell-media boundary, at $x = Ih$, assume the nanoparticle is removed from the system (associated with a cell) with probability $P_{\text{assoc}}h$ if it attempts to transition through the cell-media boundary. The transition probabilities are therefore

$$(x(t+\tau), y(t+\tau), z(t+\tau)) = \begin{cases} (Ih, y(t), z(t)) & \text{with probability} \\ & 1 - 6D\tau/h^2 + (1 - P_{\text{assoc}}M_{j,k}h)\left(\frac{C^{\max} - C_{j,k}}{C^{\max}}\right)\left(\frac{D\tau}{h^2} + \frac{V\tau}{2h}\right), \\ (x(t)-h, y(t), z(t)) & \text{with probability } D\tau/h^2 - V\tau/2h, \\ (x(t), y(t)+h, z(t)) & \text{with probability } D\tau/h^2, \\ (x(t), y(t)-h, z(t)) & \text{with probability } D\tau/h^2, \\ (x(t), y(t), z(t)+h) & \text{with probability } D\tau/h^2, \\ (x(t), y(t), z(t)-h) & \text{with probability } D\tau/h^2, \end{cases}$$

and removed from the system with probability $P_{\text{assoc}}M_{j,k}h(D\tau/h^2 + V\tau/2h)(C^{\max} - C_{j,k})/C^{\max}$.

Consider now the evolution of the number of nanoparticles in the voxel $(i, j, k)$

$$
\begin{aligned}
N_{(i,j,k)}(t+\tau) = {} & (1 - \frac{6D\tau}{h^2})N_{(i,j,k)}(t) \\
& + \frac{D\tau}{h^2}\Big(N_{(i+1,j,k)}(t) + N_{(i-1,j,k)}(t) + N_{(i,j+1,k)}(t) + N_{(i,j-1,k)}(t) + N_{(i,j,k+1)}(t) + N_{(i,j,k-1)}(t)\Big) \\
& + \frac{V\tau}{2h}\Big(N_{(i-1,j,k)}(t) - N_{(i+1,j,k)}(t)\Big).
\end{aligned}
$$

Rearranging, we obtain

$$
\begin{aligned}
\frac{N_{(i,j,k)}(t+\tau) - N_{(i,j,k)}(t)}{\tau} = {} & \frac{D}{h^2}\Big(N_{(i+1,j,k)}(t) + N_{(i-1,j,k)}(t) + N_{(i,j+1,k)}(t) \\
& + N_{(i,j-1,k)}(t) + N_{(i,j,k+1)}(t) + N_{(i,j,k-1)}(t) - 6N_{(i,j,k)}(t)\Big) \\
& + \frac{v}{2h}\Big(N_{(i-1,j,k)}(t) - N_{(i+1,j,k)}(t)\Big).
\end{aligned}
$$

Taking the limit $\tau \to 0$, $h \to 0$ [2], we recover the traditional dosage model.

For the cell-media boundary, the evolution of the number of nanoparticles in a boundary voxel is

$$
\begin{aligned}
N_{(I,j,k)}(t+\tau) = {} & \left(1 - \frac{6D\tau}{h^2} + (1 - P_{\text{assoc}}M_{j,k}h)\left(\frac{D\tau}{h^2} + \frac{V\tau}{2h}\right)\left(\frac{C^{\max} - C_{j,k}}{C^{\max}}\right)\right)N_{(I,j,k)}(t) \\
& + \frac{D\tau}{h^2}\Big(N_{(I-1,j,k)}(t) + N_{(I,j+1,k)}(t) + N_{(I,j-1,k)}(t) + N_{(I,j,k+1)}(t) + N_{(I,j,k-1)}(t)\Big) \\
& + \frac{V\tau}{2h}N_{(I-1,j,k)}(t).
\end{aligned}
$$

Upon rearranging, we obtain

$$
\begin{aligned}
\frac{N_{(I,j,k)}(t+\tau) - N_{(I,j,k)}(t)}{\tau} = {} & \frac{D}{h^2}\Big(N_{(I,j+1,k)}(t) + N_{(I,j-1,k)}(t) + N_{(I,j,k+1)}(t) + N_{(I,j,k-1)}(t) - 4N_{(I,j,k)}(t)\Big) \\
& + \frac{V}{2h}\left(N_{(I-1,j,k)}(t) + N_{(I,j,k)}(t) - P_{\text{assoc}}M_{j,k}h\left(\frac{C^{\max} - C_{j,k}}{C^{\max}}\right)N_{(I,j,k)}(t)\right) \\
& + \frac{D}{h^2}\left(N_{(I-1,j,k)}(t) - N_{(I,j,k)}(t) - P_{\text{assoc}}M_{j,k}h\left(\frac{C^{\max} - C_{j,k}}{C^{\max}}\right)N_{(I,j,k)}(t)\right).
\end{aligned}
$$

This is equivalent to

$$
\begin{aligned}
\sqrt{\tau}\frac{N_{(I,j,k)}(t+\tau) - N_{(I,j,k)}(t)}{\tau} = {} & D\sqrt{\tau}\left(\frac{N_{(I,j+1,k)}(t) + N_{(I,j-1,k)}(t) + N_{(I,j,k+1)}(t) + N_{(I,j,k-1)}(t) - 4N_{(I,j,k)}(t)}{h^2}\right) \\
& + \frac{V\sqrt{\tau}}{2h}\left(N_{(I-1,j,k)}(t) + N_{(I,j,k)}(t) - P_{\text{assoc}}M_{j,k}h\left(\frac{C^{\max} - C_{j,k}}{C^{\max}}\right)N_{(I,j,k)}(t)\right) \\
& + \frac{D\sqrt{\tau}}{h}\left(\frac{N_{(I-1,j,k)}(t) - N_{(I,j,k)}(t)}{h} - P_{\text{assoc}}M_{j,k}\left(\frac{C^{\max} - C_{j,k}}{C^{\max}}\right)N_{(I,j,k)}(t)\right).
\end{aligned}
$$

Taking the limit $\tau \to 0$, $h \to 0$ such that $\sqrt{\tau}/h$ is finite [2], we obtain

$$
D\frac{\partial N(x,y,z,t)}{\partial x} - VN(x,y,z,t) = DP_{\text{assoc}}M_{j,k}\left(\frac{C^{\max} - C_{j,k}}{C^{\max}}\right)N(x,y,z,t), \text{ at } x = L.
$$

This is consistent with the boundary condition in the traditional dosage model provided

$$
P_{\text{assoc}} = \alpha/D.
$$

| Parameter | Definition | 1032nm Capsule-RAW | 282nm Coreshell-HeLa | 150nm Coreshell-RAW |
|---|---|---|---|---|
| $k_b$ | Boltzmann constant ($J \cdot K^{-1}$) | $1.38 \cdot 10^{-23}$ | $1.38 \cdot 10^{-23}$ | $1.38 \cdot 10^{-23}$ |
| $T$ | Temperature (K) | 310.15 | 310.15 | 310.15 |
| $\eta$ | Dynamic viscosity (Pa·s) | $10^{-3}$ | $10^{-3}$ | $10^{-3}$ |
| $d$ | Nanoparticle diameter (m) | $1.32 \cdot 10^{-6}$ | $2.82 \cdot 10^{-7}$ | $1.5 \cdot 10^{-7}$ |
| $g$ | Gravitational acceleration ($m^2 \cdot s^{-1}$) | 9.81 | 9.81 | 9.81 |
| $\rho$ | Nanoparticle density ($kg \cdot m^{-3}$) | 1009 | 1850 | $1.25 \cdot 10^4$ |
| $\rho_m$ | Fluid density ($kg \cdot m^{-3}$) | 1000 | 1000 | 1000 |
| $S$ | Cell confluence | 0.21 | 0.41 | 0.21 |
| $P_0$ | Nanoparticle concentration ($m^{-3}$) | $9.95 \cdot 10^{12}$ | $9.95 \cdot 10^{12}$ | $9.95 \cdot 10^{12}$ |
| $C^{\max}$ (Mean) | Carrying capacity | 60 | 8 | 20 |
| $\alpha$ (Mean) | Nanoparticle-cell affinity ($m \cdot s^{-1}$) | $2.97 \cdot 10^{-9}$ | $8.37 \cdot 10^{-10}$ | $1.47 \cdot 10^{-9}$ |
| $L$ | Media depth (m) | $2.6 \cdot 10^{-3}$ | $2.6 \cdot 10^{-3}$ | $2.6 \cdot 10^{-3}$ |
| $h$ | Voxel size (m) | $2.88 \cdot 10^{-5}$ | $4 \cdot 10^{-5}$ | $2.88 \cdot 10^{-5}$ |
| $I$ | Number of voxels (x) | 91 | 66 | 91 |
| $J$ | Number of voxels (y) | 100 | 100 | 100 |
| $K$ | Number of voxels (z) | 100 | 100 | 100 |
| $1/r_{G1}$ | G1 phase length (s) | $2.97 \cdot 10^4$ | $2.4 \cdot 10^4$ | $2.97 \cdot 10^4$ |
| $1/r_S$ | S phase length (s) | $1.89 \cdot 10^4$ | $3.0 \cdot 10^4$ | $1.89 \cdot 10^4$ |
| $1/r_{G2}$ | G2 phase length (s) | $4.05 \cdot 10^3$ | $1.05 \cdot 10^4$ | $4.05 \cdot 10^3$ |
| $1/r_M$ | M phase length (s) | $1.35 \cdot 10^3$ | $3.0 \cdot 10^3$ | $1.35 \cdot 10^3$ |
| $\tau$ | Gillespie timestep (s) | 10 | 10 | 2 |
| $\Delta t$ | Euler timestep (s) | 1 | 1 | 1 |
| $\Delta x$ | Node spacing (m) | $2.62 \cdot 10^{-7}$ | $2.62 \cdot 10^{-7}$ | $2.62 \cdot 10^{-7}$ |

Table 1: Table of parameters for both the hybrid dosage model and voxel-based model. Definitions of individual parameters are as per the text. Parameters $T$, $\eta$, $d$, $\rho$, $\rho_m$, $S$, $P_0$, $C_{\max}$, $\alpha$, $L$ are obtained from [3]. Parameters $r_{G1}$, $r_S$, $r_{G2}$, $r_M$ are estimated from [4] for the 282nm Coreshell-HeLa experiments and from [8] for the 1032nm Capsule-RAW and 150nm Coreshell-RAW experiments. All other parameters are well-established or are numerical solver (user-selected) parameters.

### Model Parameters

The parameters used to obtain numerical solutions to the hybrid dosage model and the voxel-based model are presented in Table 1 for each experiment. Note that for These parameters are used in all figures in the main document, with the exception of Figure 7, where the nanoparticle-cell affinity and cell carrying capacity are obtained via nonlinear least squares. For Figure 7, the nanoparticle-cell affinity parameters are $\alpha = 3.79 \cdot 10^{-8}$, $\alpha = 3.77 \cdot 10^{-7}$, and $\alpha = 1.48 \cdot 10^{-8}$ for the 1032nm Capsule-RAW, 282nm Coreshell-Hela and 150nm Coreshell-RAW experiments, respectively. The cell carrying capacity parameters are $C^{\max} = 9.59$, $C^{\max} = 7.69$, and $C^{\max} = 28.42$ for the 1032nm Capsule-RAW, 282nm Coreshell-Hela and 150nm Coreshell-RAW experiments, respectively.

### Additional Results

In Figure S1, we show that the dose obtained from the model is only Poisson distributed in the linear association regime (Fig. S1A-D). If the number of nanoparticles per cell approaches the carrying capacity, the predicted dosage distribution is not well described by the Poisson distribution (Fig. S1E-F). Comparing the dosage distribution obtained from the model (Fig. S1A-F) with the experimental data (Fig. S1G-L), we observe that the experimental data is overdisperse compared to the model predictions.

In Figure S2 we demonstrate that the number of nanoparticles associated with cells is consistent with the results obtained from traditional dosage model.

In Figure S3 we demonstrate that the apparent heterogeneity obtained from the hybrid model matches the apparent heterogeneity obtained from the voxel-based model.

In Figures S4-S8, we present the difference between the Poisson distribution (Figure S4) or Poisson-lognormal distribution (Figure S5-S8) and the dosage distribution obtained from the hybrid model. The

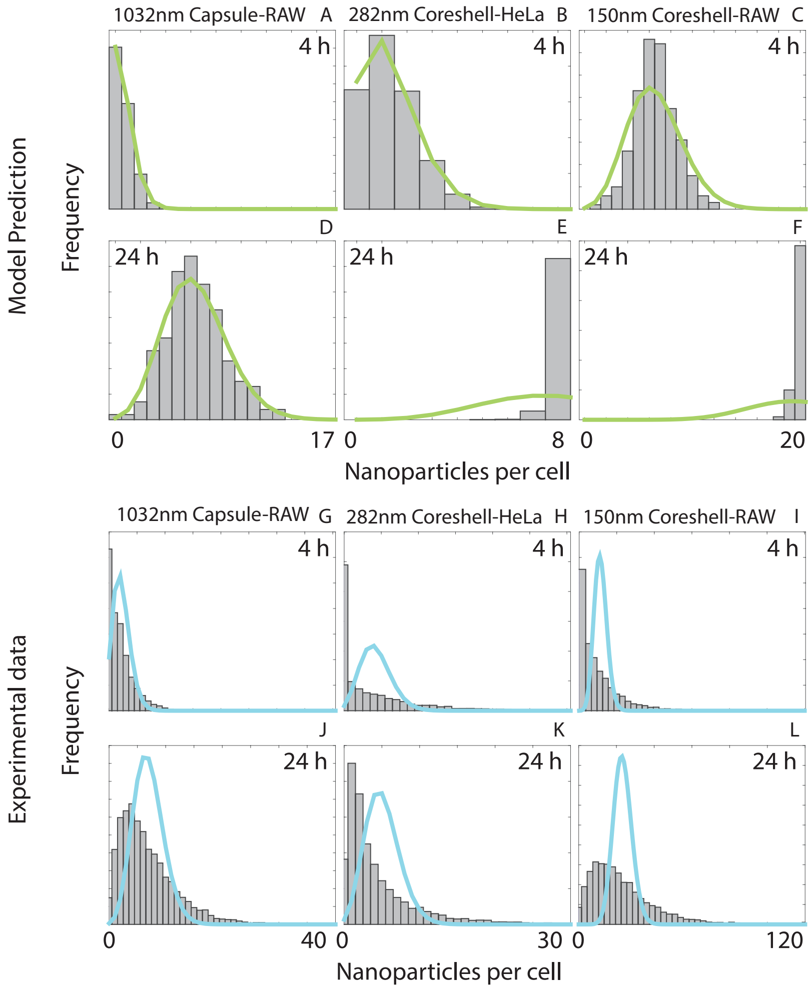


Figure S1: **Stochastic motion does not account for all variation in dosage distributions.** Histograms of the number of associated nanoparticles per cell for three different nanoparticle-cell pairs after (**A-C**) 4 hours and (**D-F**) 24 hours obtained from the voxel-based modelling framework, and after (**G-I**) 4 hours and **J-L.** 24 hours from the experimental data. The (**A-F**) green and (**G-L**) cyan lines correspond to the Poisson distribution that best fits the dosage distribution.

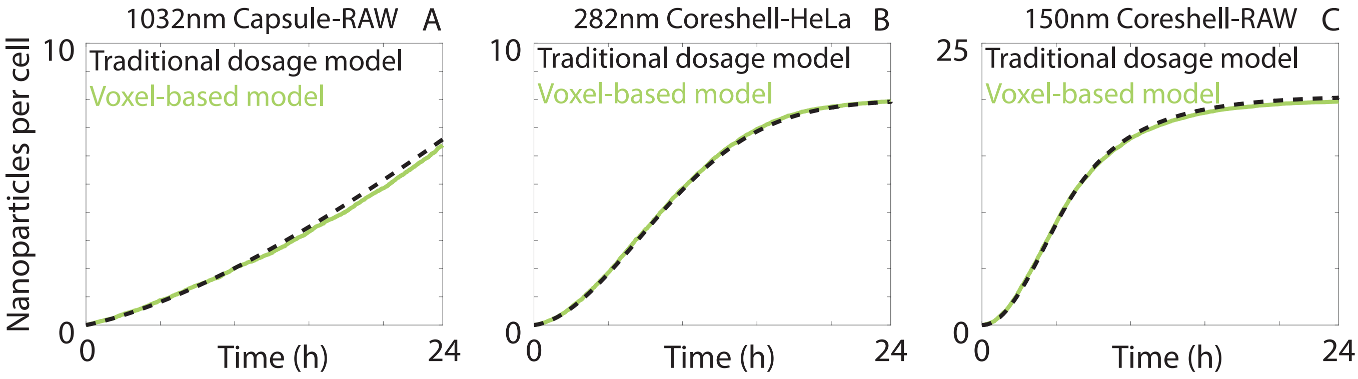


Figure S2: **Comparison in mean nanoparticle dose between the traditional dosage model and the voxel-based model.**

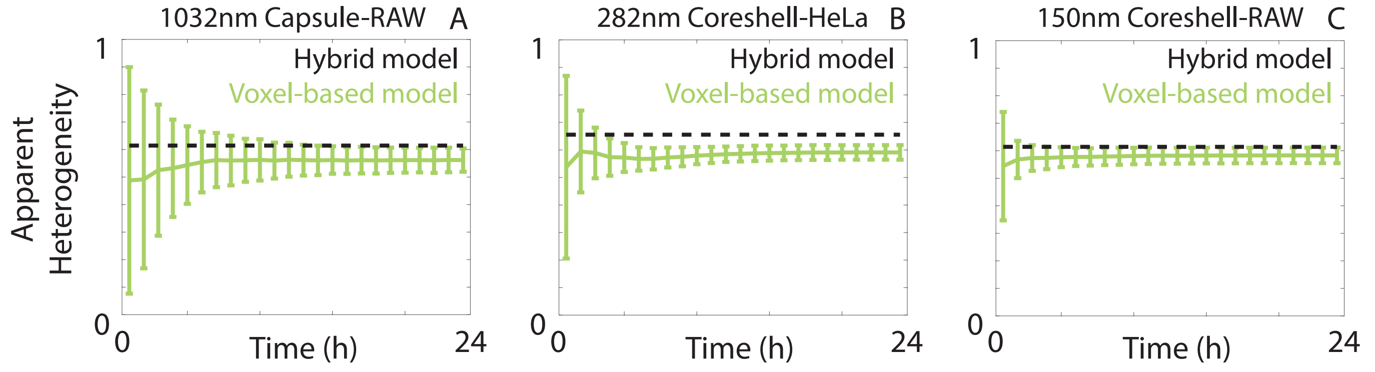


Figure S3: **Comparison in apparent heterogeneity between the hybrid model and two hundred realisations of the voxel-based model.** Error bars correspond to one standard deviation.

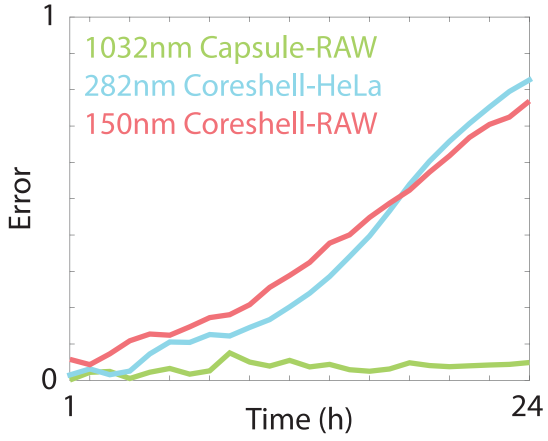


Figure S4: **Difference between the Poisson distribution and the model dosage distribution in Figure S1.**

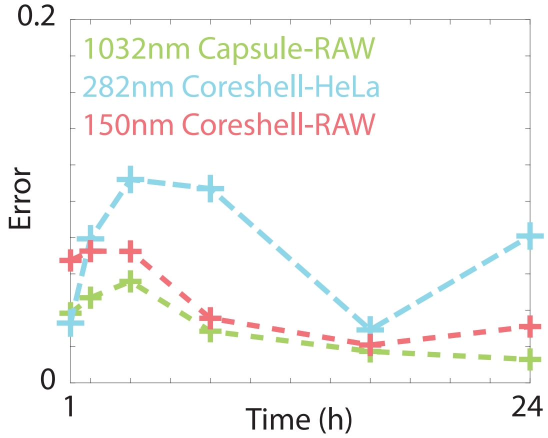


Figure S5: **Difference between the Poisson-lognormal distribution and the experimental dosage distribution in Figure 3 (main document).**

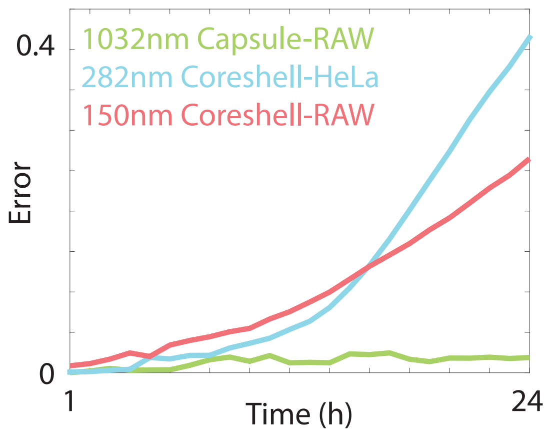


Figure S6: **Difference between the Poisson-lognormal distribution and the model dosage distribution in Figures 4D-F (main document).**

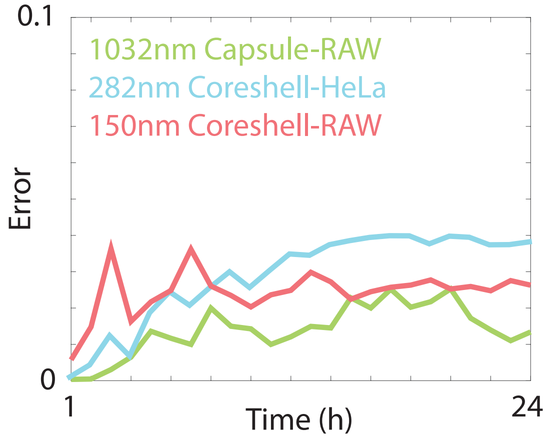


Figure S7: **Difference between the Poisson-lognormal distribution and the model dosage distribution in Figures 4J-L (main document).**

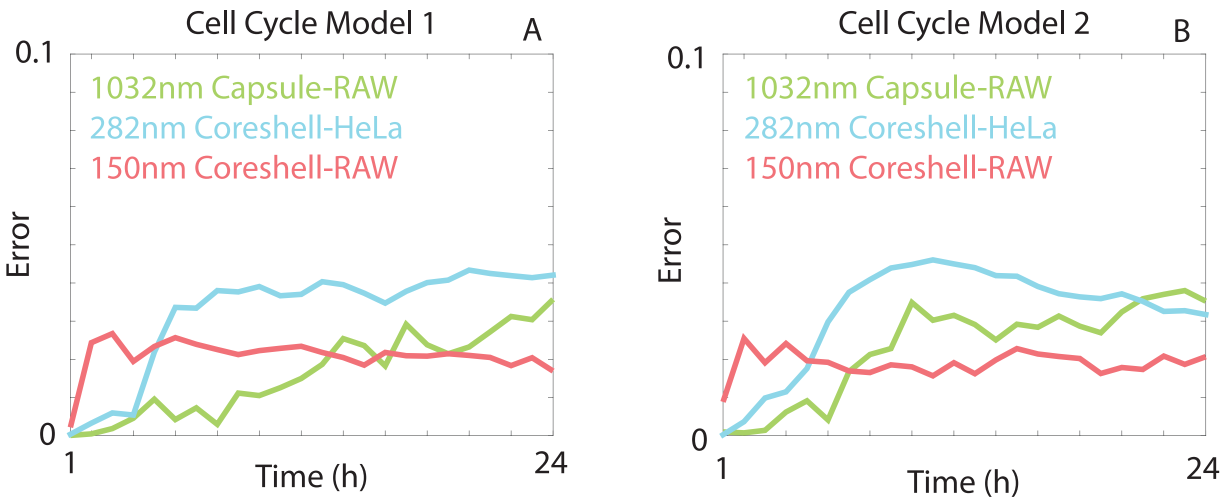


Figure S8: **Difference between the Poisson-lognormal distribution and the model dosage distribution in Figure 5 (main document) for (A) Cell Cycle Model 1 and (B) Cell Cycle Model 2.**

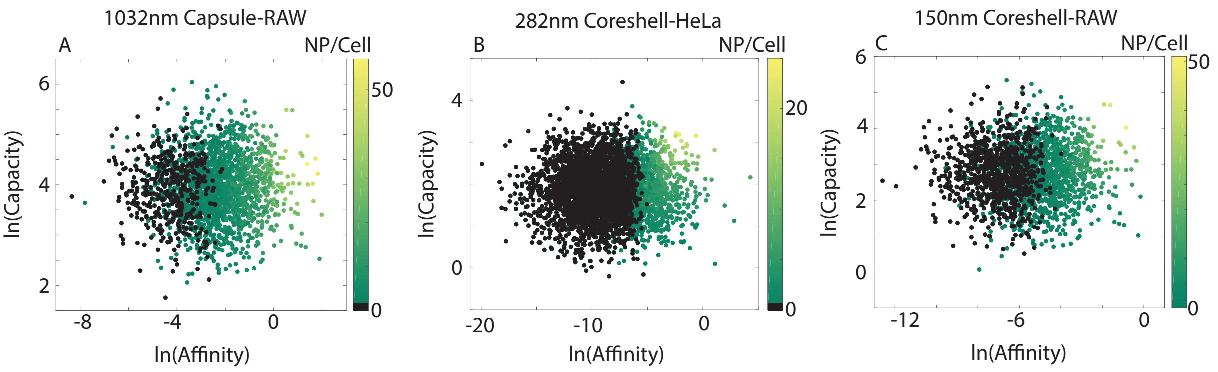


Figure S9: **Scatter plots highlighting the relationship between nanoparticle-cell affinity, cell carrying capacity and the number of associated nanoparticles**. Black circles correspond to cells with nanoparticle-cell affinity and cell carrying capacity values that resulted in zero associated nanoparticles.

difference is defined as

$$\text{Error} = \frac{1}{N_{\text{cells}}}\left(\sum_{m=0}^{\max(C_{j,k})}\left(N_{\text{cells}}p(m) - h(m)\right)^2\right)^{1/2},$$

where $N_{\text{cells}}$ is the number of cells in the simulation, $p(m)$ is the probability density function for the appropriate distribution and $h(m)$ is the number of cells that are associated with $m$ nanoparticles. Error values that grow with time indicate that the assumptions regarding the distribution are inappropriate. As expected, the error in Figure S4 and Figure S6 grows for the 282nm Capsule-HeLa and 150nm Capsule-RAW experiments, due to the constant carrying capacity. For all other experiments we observe that error is consistently low and hence the assumption of the Poisson-lognormal distribution appears appropriate.

In Figure S9 we present scatter plots of nanoparticle-cell affinity values and carrying capacity values, as well as the corresponding number of associated nanoparticles per cell at the end of 24 hours, as predicted by the voxel-based model. Parameter values that correspond to cells that do not associate with any nanoparticles are coloured in black. We observe that such cells typically have lower nanoparticle-cell affinity values and low cell carrying capacity values, which would impose a selective pressure towards these values if the nanoparticles contained a therapeutic agent.
